# Characterization of a nanobody-epitope tag interaction and its application for receptor engineering

**DOI:** 10.1101/2022.05.08.491096

**Authors:** Chino C. Cabalteja, Shivani Sachdev, Ross W. Cheloha

**Affiliations:** Laboratory of Bioorganic Chemistry; National Institute of Diabetes, Digestive, and Kidney Disease; National Institutes of Health

## Abstract

Peptide epitope tags offer a valuable means for detection and manipulation of protein targets for which high quality detection reagents are not available. Most commonly used epitope tags are bound by conventional, full-size antibodies (Abs). The complex architecture of Abs complicates their application in protein engineering and intracellular applications. To address these shortcomings, single domain antibodies (nanobodies, Nbs) that recognize short peptide epitopes have become increasingly prized. Here we characterize the interaction between a Nb (Nb_6E_) and a 14-mer peptide epitope. We identify residues in the peptide epitope essential for high affinity binding. Using this information in combination with computational modeling we propose a mode of interaction between Nb_6E_ and this epitope. We apply this nanobody-epitope pair to augment the potency of a ligand at an engineered adenosine A2A receptor. This characterization of the nanobody-epitope pair opens the door to diverse applications including mechanistic studies of G protein-coupled receptor function.

## Introduction

Antibodies (Abs) are prized as reagents for the highly specific recognition of biological molecules of interest, especially for protein targets. Abs are widely used in basic biomedical research for the detection and manipulation of proteins and in therapeutic applications for blocking the action of soluble proteins in circulation or receptors found on the cell surface. In these settings, the identification of antibodies with high affinity, specificity, and stability can propel research and therapeutic progress^1^. For some types of targets the identification of antibodies with such desirable properties is difficult. For example, many members of the G protein-coupled receptor (GPCR) superfamily are mostly found embedded in the plasma membrane and have little extracellular surface area exposed for binding by Abs^2^. In such cases, it is often useful to tag proteins of interest with a short peptide epitope tag to allow for recognition by high quality and well-established anti-epitope tag Abs^3^. Abs that bind to epitope tags such as myc, HA, and FLAG have been used routinely for decades as immunological detection reagents.

Conventional Abs are comprised of four polypeptide chains (two heavy chains and two light chains) that require glycosylation and disulfide bond formation for assembly, folding, and function. This complex architecture makes the expression of antibodies in the reducing environment of the cellular cytoplasm difficult or impossible. Attempts to create fusions between Abs and bioactive proteins of interest often results in reduced production yields, a loss in folded protein stability, or undesirable proteolytic cleavage^4^. To circumvent some of these problems, Ab fragments that maintain the high affinity and specificity of full-size immunoglobulins have been developed. Fragments derived directly from conventional Abs include constructs known as single chain-fragment variable (scFv) and fragment antigen-binding (Fab); however, these constructs still possess some of the limitations seen in full-size Abs^5^. For example, scFvs are commonly used as recognition agents in chimeric antigen receptor T cell (CAR-T) constructs; however, scFv misfolding or mispairing often leads to receptor aggregation or other complications^6^. Single domain antibodies derived from the variable region of camelid heavy chain only antibodies (VHHs or nanobodies, Nbs) offer an alternative to address some of these issues. Nbs are the smallest antibody fragment that maintains full affinity and they usually do not require disulfide bond formation or glycosylation for function^7^.

Efforts have been undertaken to identify Nbs that bind to peptide epitope tags^7^. Nbs that bound to the human immunodeficiency virus protein gp41 were shown to bind to short peptide fragments of this protein, which could be applied as epitope tags (MoonTag, PepTag)^8,9^. Alternatively, a collection of Nbs specifically raised against a prototypical, synthetic α-helical peptide (Alfa Tag) have been used for a variety of applications^10,11^. Nanobodies raised against the mammalian proteins α-synuclein^12^, β-catenin^13^, CXCR2 (ref. 14,15), and UBC6e^16^ have been shown to bind to small peptide epitopes taken from these proteins. Previous work has shown that a nanobody (previously called VHH05 or VHH_6E_, here named Nb_6E_) binds to a 14-mer peptide derived from the protein UBC6e (6E tag) with low nM affinity^16^. Subsequent work showed that the 6E tag could be appended via genetic fusion to the extracellular portion of parathyroid hormone receptor-1 (PTHR1), a GPCR, where it was served effectively as an epitope tag without sacrificing receptor function^17,18^. Success with the Nb_6E_-6E tag pair spurred us to further characterize this bimolecular interaction. Herein we provide a structure-activity relationship study for the Nb_6E_-6E tag interaction, generate a computational model to contextualize these experimental data, and apply this nanobody-tag pair to interrogate GPCR signaling.

## Results and discussion

We first sought to identify positions within the 6E tag peptide that are important for interaction with Nb_6E_. Using solid phase peptide synthesis, we prepared a comprehensive scan library where residues at each position were swapped with Ala except for naturally occurring Ala residues, which were swapped with Glu (Figure 1, Supporting Figure 1 for MS). All peptides also contain a triglycine motif at the N-terminus for use in Sortase A-mediated labeling^19^. Based on preliminary data, we also prepared an analogue containing the non-natural amino acid aminoisobutyric acid (Aib), which promotes helix formation. We then evaluated the binding of these peptides with Nb_6E_ using a variety of methods.

**Figure 1.**
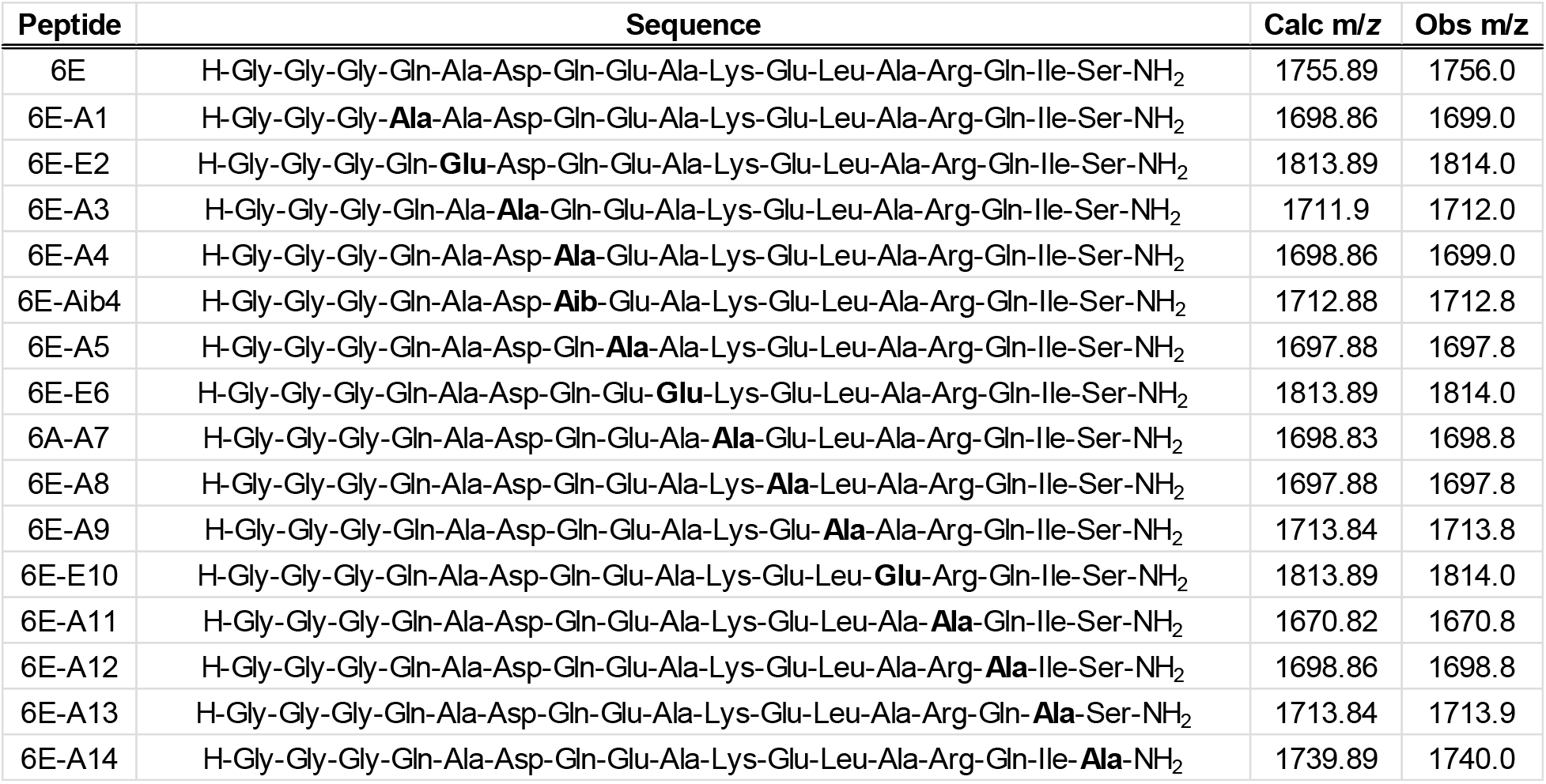
Sequences of peptides used in this study. “Obs m/z” represents the mass corresponding to the most abundant ion recorded in characterization using liquid chromatography/mass spectrometry (Supporting Figure 1).

We first developed a competition-based enzyme-linked immunosorbent assay (ELISA) in which each member of the scan library was tested for its ability to inhibit binding of Nb_6E_ to immobilized 6E peptide. To facilitate efficient immobilization on the surface of the ELISA plate for 6E peptide we used sortase-mediated labeling (sortagging) to link 6E with enhanced green fluorescent protein (Figure 2A, Supporting Figure 1). We also labeled Nb_6E_ with biotin using sortagging to permit detection with streptavidin-horse radish peroxidase (SA-HRP) conjugate. Several peptides from the scan library performed similarly to the prototype peptide in this assay (Figure 2B, Supporting Figure 2), indicating maintenance of binding affinity upon Ala substitution. Other members of the scan library exhibited modestly weakened binding. A few analogues (6E-E2, A3, E6, A9, A10) exhibited dramatically reduced binding, suggesting these residues are important for Nb_6E_ association.

**Figure 2.**
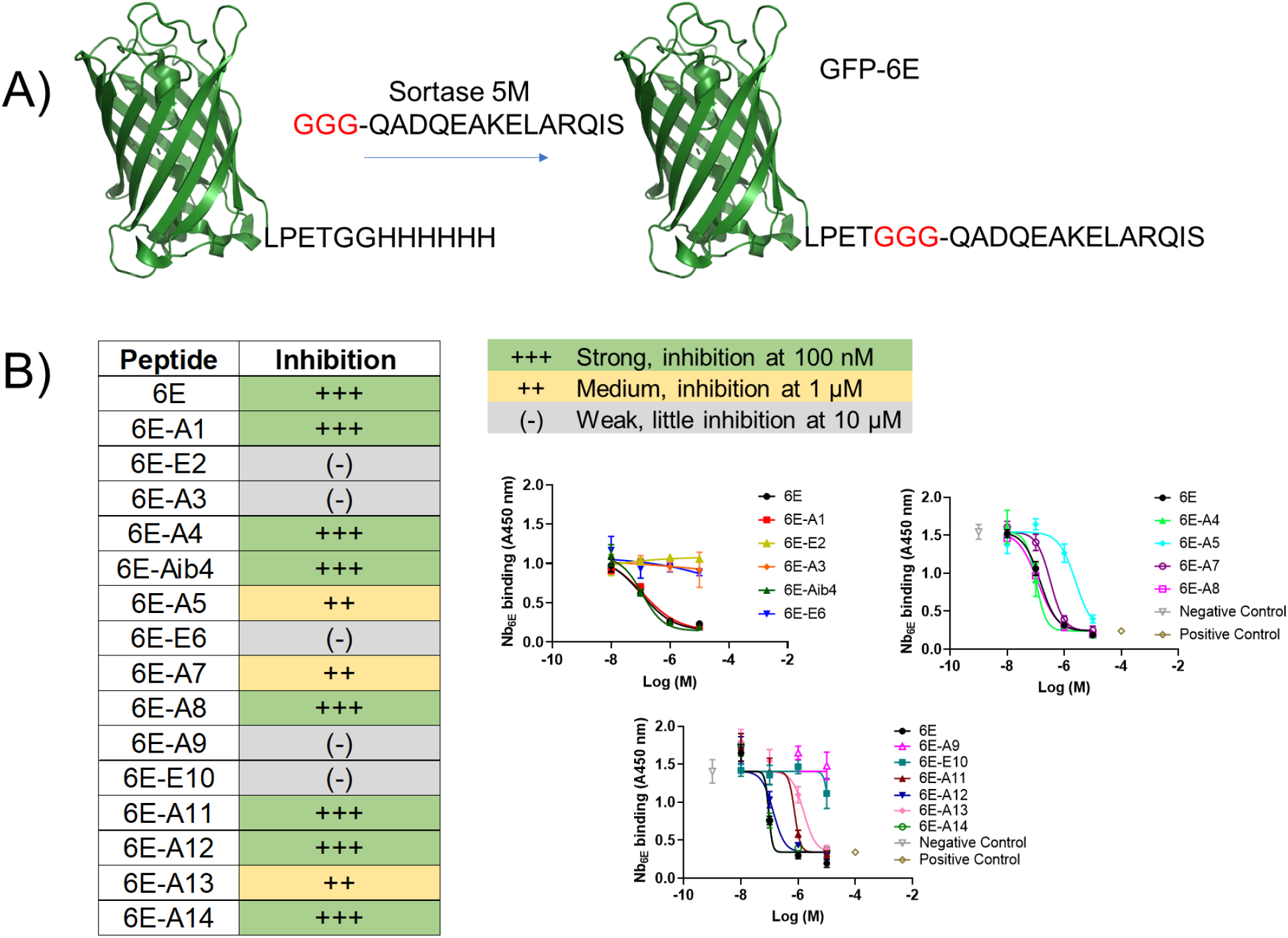
Characterization of 6E peptide-Nb_6E_ binding by competition ELISA. A) Schematic for the preparation of GFP-6E conjugates using sortagging for immobilization in ELISA. See methods for reaction details. B) Summary data for analysis of peptide analogue binding using ELISA. Peptides were categorized into three groups based on their performance in these assays. Representative data from individual assays are shown at right. Data points show mean ± SD from technical replicates in a single experiment. Positive control data points correspond to signal measured in the absence of Nb_6E_-biotin. Negative control data points correspond to signal observed in the absence of competitor peptide (but with Nb_6E_-biotin). Curves result from fitting a four-parameter sigmoidal dose-response model to the data (except for weak (−) compounds, which show connected points). Independent replicate data are shown in Supporting Figure 2. Each analogue was tested in ≥ 3 independent experiments. If peptide inhibition categories varied between independent experiments, we categorized compounds in this table based on their modal performance in assays.

To further characterize binding for selected analogues we performed surface plasmon resonance (SPR) experiments (Figure 3). We chose representative peptides from strong, medium, and weak binder categories for these experiments. We immobilized biotinylated Nb_6E_ onto sensor chips coated with streptavidin (SA) and flowed peptides over the coated sensor chips. All peptides demonstrated binding and dissociation in SPR sensorgrams, although the data was not well fit with a simple one-state binding model, requiring the use of a more complex two-state model (see Supporting Methods). The data obtained in SPR experiments mirrored observations from ELISA. 6E peptide bound with the highest affinity, 6E-A5 with modestly reduced affinity, and 6E-A9 with the lowest affinity (Figure 3A-B). Binding association and dissociation parameters were variable among the different analogues tested, with interpretation complicated by use of the two-state model (Supporting Figure 3).

**Figure 3.**
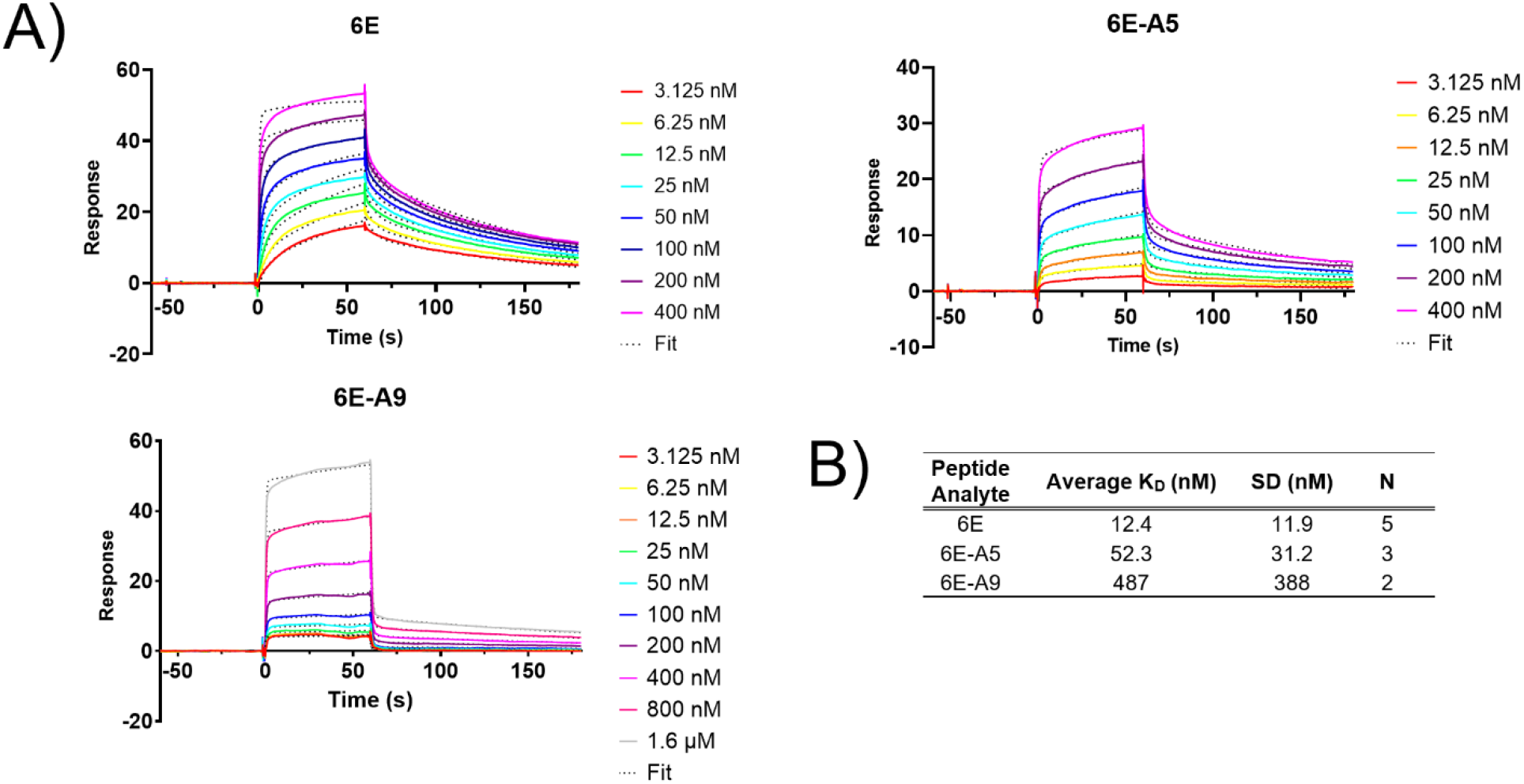
Characterization of Nb_6E_-6E peptide association and dissociation by surface plasmon resonance assays. A) Representative sensorgrams showing binding and dissociation of 6E peptides to immobilized Nb_6E_. Nb_6E_ was conjugated to biotin at its C-terminus using sortagging and immobilized onto sensor chips functionalized with streptavidin. Peptides (at indicated concentrations) were flowed over functionalized sensor chips for 60 s (association) followed by 120 s of buffer flow with no peptide (dissociation). Data were fit a two-state binding model with local R-max with curve fits shown as dotted lines. B) Tabulation of binding parameters from independent replicate experiments. K_D_ averages (mean) and SD were calculated using K_D_ values derived for the indicated number of independent experiments (“N”).

The binding properties of 6E analogues were also tested in a biological milieu. We developed a flow cytometry-based assay using HEK293 cells stably transfected to express the adenosine A2A receptor (a GPCR) fused at its N-terminus with Nb_6E_ [named A2AR(Nb_6E_)]. This receptor topology places Nb_6E_ at the extracellular face of the plasma membrane, allowing for assessment of binding on live cells expressing this receptor (Figure 4A). We performed a competition assay in which a fluorescein tagged version of 6E peptide (FAM-6E-C14, tracer peptide) was mixed with varying concentrations of unlabeled analogues of 6E (6E, 6E-Aib4, 6E-A5, 6E-A9) and added to cells (Figure 4B-D, Supporting Figure 4). The extent of binding was quantified through use of an alexafluor-647 (AF647) tagged anti-fluorescein secondary antibody, with analysis of labeled cells by flow cytometry. As expected, cells expressing A2AR(Nb_6E_) showed intense staining with tracer peptide. Each of the unlabeled peptides inhibited binding of tracer peptide to A2AR(Nb_6E_) in a concentration-dependent manner. Binding was quantified by measurement of median fluorescence intensity (MFI) in the AF647 channel (Figure 4C-D, Supporting Figure 5). The trends observed mirrored those in previous assays, with 6E and 6E-Aib4 showing strong inhibition of tracer peptide binding, 6E-A5 modest inhibition, and 6E-A9 weak inhibition. Results from flow cytometry experiments were consistent with fluorescence microscopy analysis (Supporting Figure 6).

**Figure 4.**
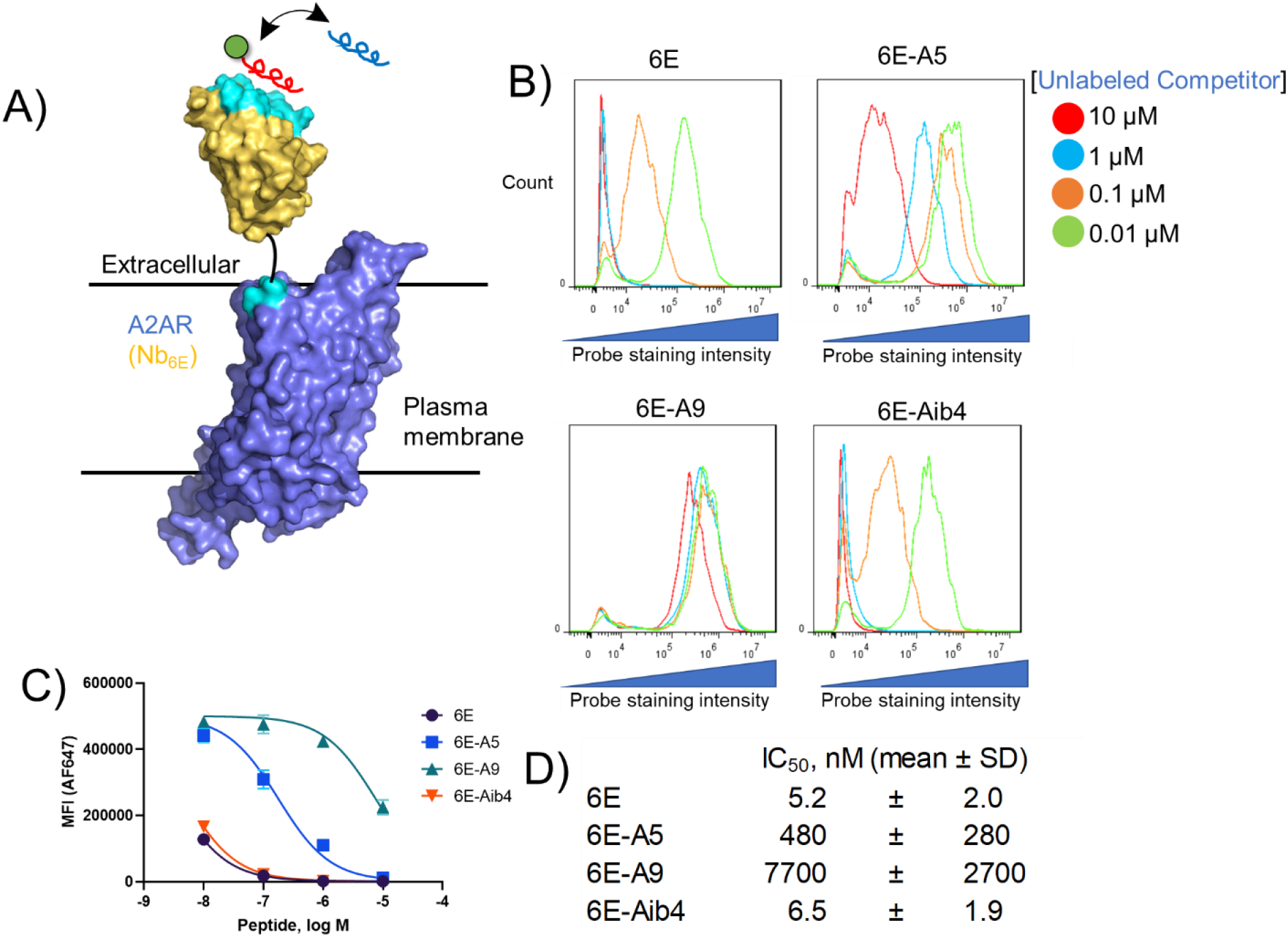
Assessment of Nb_6E_-6E peptide interaction on the surface of mammalian cells. A) Schematic of a competition binding assay involving cells expressing A2AR(Nb_6E_), a fluorescein-labeled tracer peptide (red), and unlabeled competitor peptides (blue). B) Representative data for flow cytometry analysis of the inhibition of fluorescein-labelled tracer peptide binding by indicated concentrations of unlabeled competitor peptides. Tracer peptide (10 nM) and unlabeled competitor peptides were incubated with cells expressing A2AR(Nb_6E_), followed by washing, and detection with anti-fluorescein secondary antibody labeled with AF647. Data is presented as a histogram of the intensity of AF647 staining of live cells. Data from independent replicate experiments are shown in Supporting Figure 4. C) Median fluorescence intensity (MFI) quantification of the histograms presented in panel 4B. Curves correspond to application of a sigmoidal dose-response response model to the data. Data from independent replicate experiments are shown in Supporting Figure 5. D) Mean IC_50_ values for inhibition of FAM-6E-C14 binding for each unlabeled peptide tested. IC_50_ values are reported as mean ± SD from three independent experiments.

Based on these binding data we sought to develop a model to describe the interaction between Nb_6E_ and 6E peptide. Efforts to characterize the Nb_6E_-6E complex by crystallography failed to yield suitable crystals (data not shown). Recent advances in protein structure prediction methods have enabled rapid and accurate predictions of protein structure^20,21^. Since high resolution structural information for UBC6e is currently not available, we elected to model the structure of full-length human UBC6e using Alphafold2. The output of our query suggested that the 6E epitope region adopts an α-helical conformation that makes contact with the folded core of the protein (Figure 5A). Alphafold2 has also been used previously to model peptide-protein complex structure^22^. Using a similar approach, we generated a model of the Nb_6E_-6E interaction in which the 6E peptide forms a helical structure docked near portions of Nb_6E_ that include the complementarity determining regions (CDRs) (Figure 5B-D), in line with previously published nanobody-peptide epitope crystal structures^10,12,13^. This model of binding is supported by structure-activity relationship data above, in which substitutions at residues predicted to occur at the binding interface with Nb_6E_ cause substantial losses in binding (Figure 1, Figure 2B). Further evidence for a helical binding conformation for 6E comes from the observation that the incorporation of a non-natural amino acid that promotes helix formation (Aib, Figure 1)^23^ supports high affinity binding.

**Figure 5.**
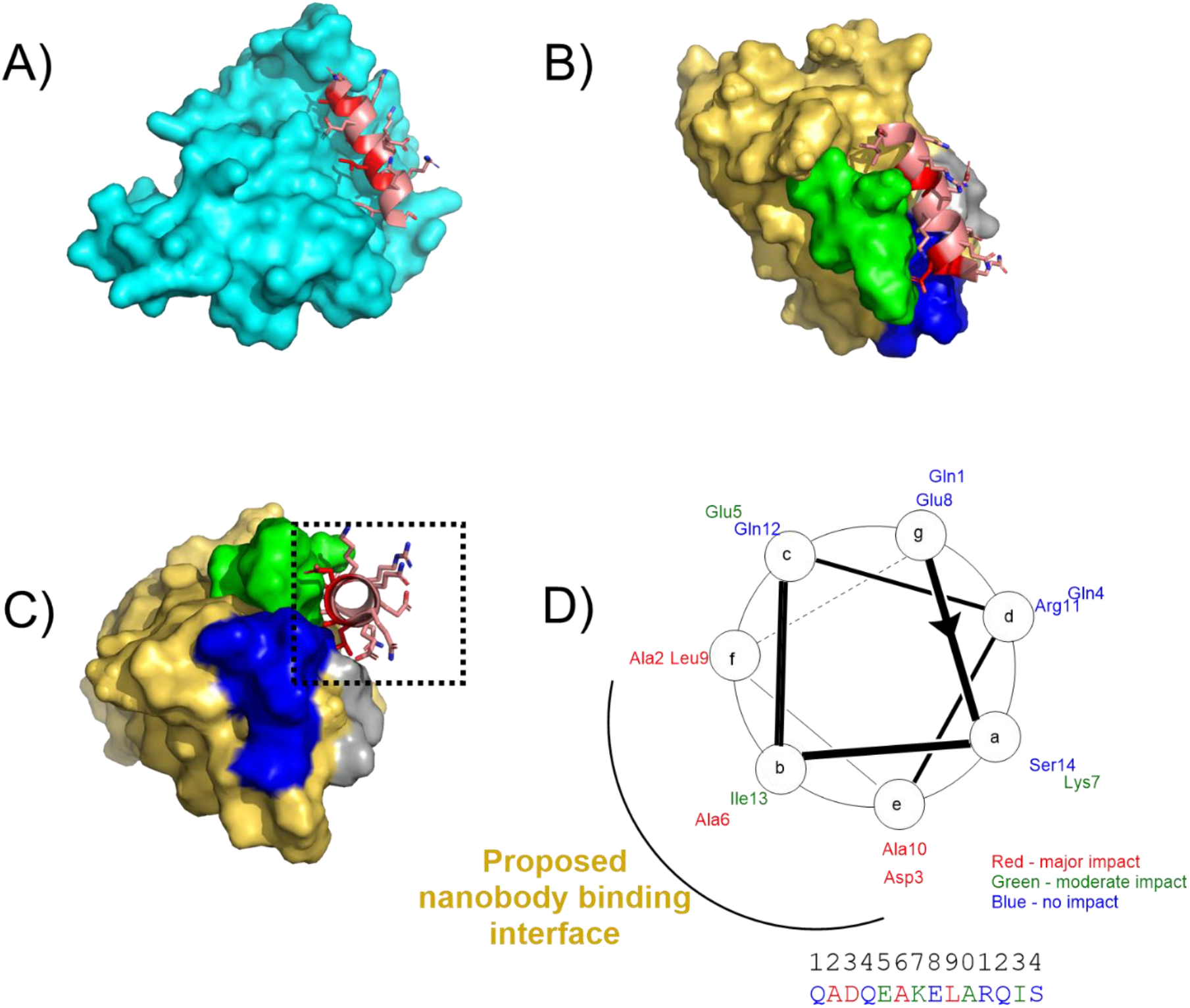
Models of 6E peptide structure and interactions. A) Model of the 6E peptide portion of UBC6e (cartoon, shown in salmon with residues important for binding to Nb_6E_ in red) in the context of UBC6e (remainder of the modeled protein shown in cyan surface). The structural model was generated using Alphafold 2^20^. The structure of the C-terminal portion (residues 183 onward) of UBC6e was predicted with low confidence and is omitted from this structure. The color-coded sequence is shown in Supporting Figure 7. B) Model of 6E peptide (salmon/red cartoon) bound to Nb_6E_ (gold surface) with Nb complementary determining regions (CDRs) highlighted in contrasting colors (CDR1: gray, CDR2: green, CDR3: blue). The structure was generated by inputting a sequence consisting of Nb_6E_ fused to 6E peptide by a flexible linker peptide. The color-coded input sequence is shown in Supporting Figure 7. Note that the 6E peptide used for modeling does not contain the GGG extension at the N-terminus. C) Reoriented perspective of hypothetical complex between 6E (red/salmon) and Nb_6E_ (gold). D) Helical wheel diagram of 6E peptide with ELISA data summarized using color coding. The putative nanobody binding interface is proposed based on the structural model shown in panel C and structure-activity relationship data.

We adapted the Nb_6E_-6E epitope tag pair to engineer a chimeric A2AR ligand that binds to both the receptor orthosteric site and a high affinity, non-orthosteric binding site. First, we developed chemistry to link an A2AR agonist small molecule to 6E peptide (Figure 6A). For the A2AR agonist we used CGS21680, which has been used previously to synthesize bitopic ligands targeting A2AR and GPCR dimers^24^. The binding of CGS21680 to human A2AR has been characterized by X-ray crystallography^25^. This structure shows that the carboxylate on CGS21680 projects from the extracellular opening of A2AR, offering a logical site for linking to other ligands. To the carboxylate on CGS21680 we coupled an alkyne group connected by a short polyethylene glycol (PEG) linker. To facilitate linkage between 6E and CGS21680 we synthesized an analogue of 6E with an azide moiety appended at the peptide N-terminus (Azide-6E, Figure 6A). Using standard copper-catalyzed azide-alkyne cycloaddition (“click”) chemistry conditions^26^ we prepared a CGS21680-6E conjugate (CGS-6E, Figure 6A). We then compared CGS-6E to CGS21680 for inducing activation of A2AR(Nb_6E_) by monitoring HEK293 cells stably transfected with receptor and a cAMP-sensitive luciferase variant^27^. We found that CGS-6E was as efficacious as CGS21680 for inducing a cyclic adenosine monophosphate (cAMP) response and that it was >25-fold more potent (Figure 6B, Supporting Figure 8). This trend was not observed for cells expressing a different version of A2AR that was fused at its N-terminus with enhanced green fluorescent protein (A2AR_EGFP_, Supporting Figure 9).

**Figure 6.**
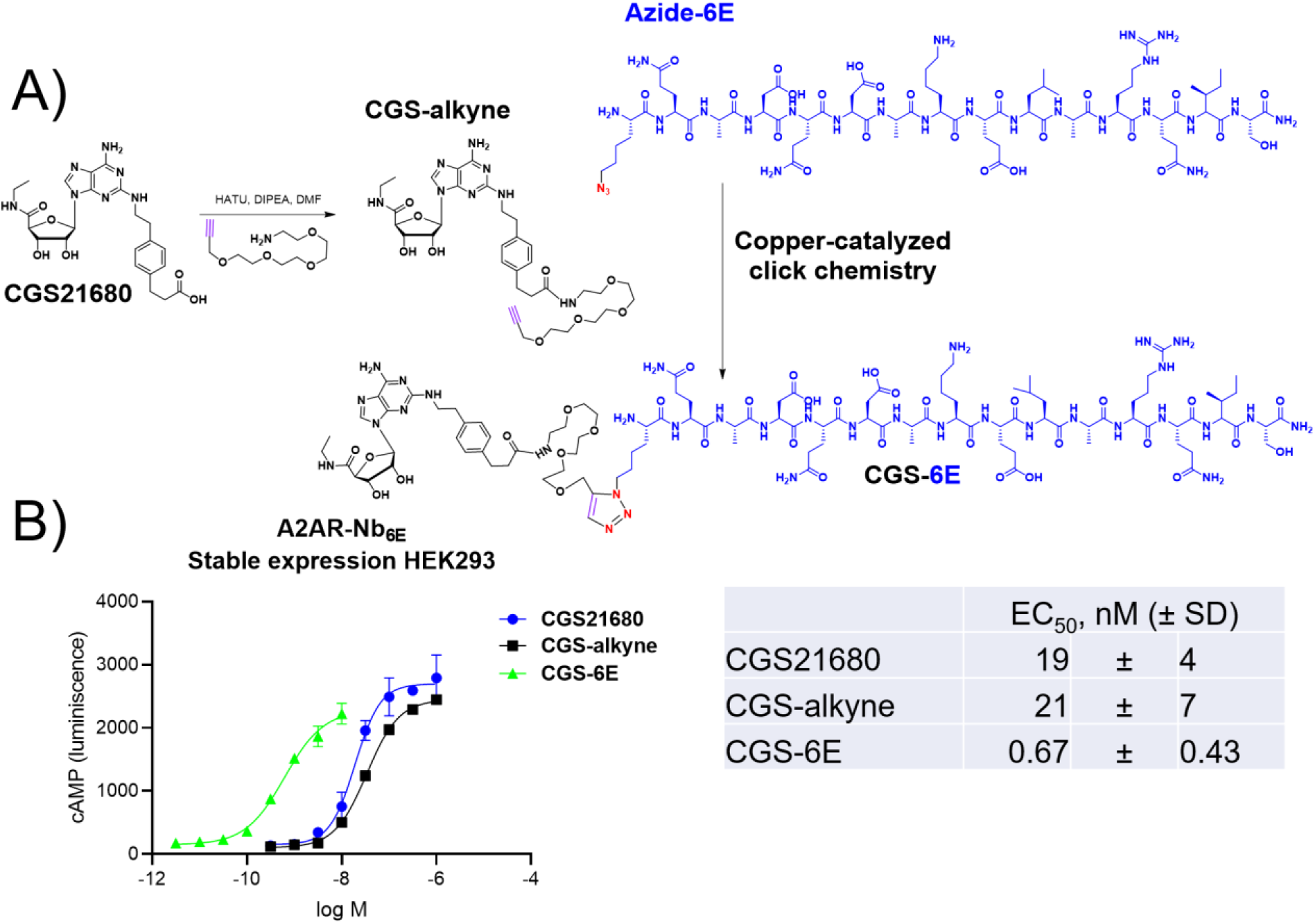
Synthesis and application of a GPCR ligand-6E peptide conjugate for enhancing ligand potency. A) Synthetic scheme for the preparation of conjugates consisting of an A2AR agonist (CGS21680) and 6E peptide using “click” chemistry. Detailed synthetic methodology and characterization is provided in the methods section. B) Evaluation of pharmacological properties of CGS-6E. (left) A representative dose-response curve for the action of indicated compounds for inducing cAMP responses on cells expressing A2AR(Nb_6E_). Data points represent mean ± SD from technical replicates in a single experiment. Additional dose-response data sets are shown in Supporting Figure 8. (right) Tabulation of compound agonist potency parameters. EC_50_ values are reported as mean ± SD from three independent experiments.

Collectively, these experiments have identified 6E peptide features important for its interaction with Nb_6E_. These data support the structural model generated here (Figure 5) for this interaction.

Given the relative paucity of nanobody-peptide epitope pairs with corresponding structure-activity relationship studies^13^, these findings should empower others to adapt these tools for studies in their system of choice. Worth noting is the use of the Nb_6E_-6E tag pair for applications as diverse as the study of Drosophila and intracellular enzyme engineering^15,28^. We demonstrate the power of these tools for studying GPCR pharmacology through engineering a receptor engrafted with an artificial epitope recognizing module (Nb_6E_), which functions to augment the potency of an engineered ligand. The engraftment of Nbs as artificial recognition domains has been previously reported for cytokine receptors^29^ and for a phosphatase^30^. More work has been extended to adapt Nbs as recognition domains for chimeric antigen receptor T cell (CAR-T) constructs for targeted immunotherapy^31,32^. This work represents the first example of the engraftment of a Nb into a GPCR, which may be useful for the design of bitopic ligands^33^. Our success in leveraging Nb-epitope interaction for improving the potency of an already high affinity ligand suggests this approach should be widely useful for basic biomedical research.

## Methods

### Synthesis

Peptides were synthesized using solid-phase synthesis with Fmoc protection of backbone amines. All compounds were purified with reverse-phase high performance liquid chromatography and compound identity was confirmed by mass spectrometry. Synthetic details are described in the supporting methods section. Mass spectrometry characterization of compounds is shown in Supporting Figure 1.

### Protein expression and labeling

Nanobodies were expressed in the *E. coli* (BL21 or WK6 strains) periplasm and purified as previously described^17^ and detailed in Supporting Methods. The sequence of Nb_6E_ has been published previously^16^. Purified nanobodies were site specifically labeled using sortagging as previously described^34^ and detailed in the Supporting Methods.

### Data calculations and display

Data were analyzed and prepared for display using GraphPad Prism, FlowJo, and Fiji (ImageJ). Flow cytometry data were quantified by measuring median fluorescence of intensity (MFI) measurements.

### Fluorescence Microscopy

HEK293 cells were analyzed as described in Supporting Methods.

### Cell culture and cell-based assays

Cell-based experiments were run with clonal cell lines derived from HEK293 (ATCC CRL-1573) stably transfected with a biosensor for cAMP^27^ and receptors of interest. Receptor sequences are found in Supporting Methods. Biological response assays were run in white walled 96-well plates and luminescence responses were recorded on a plate reader as described in Supporting Methods. Cell-based binding assays were recorded using flow cytometry as described in Supporting Methods.

