## Supporting Information for "Characterization of a nanobody-epitope tag interaction and its application for receptor engineering"

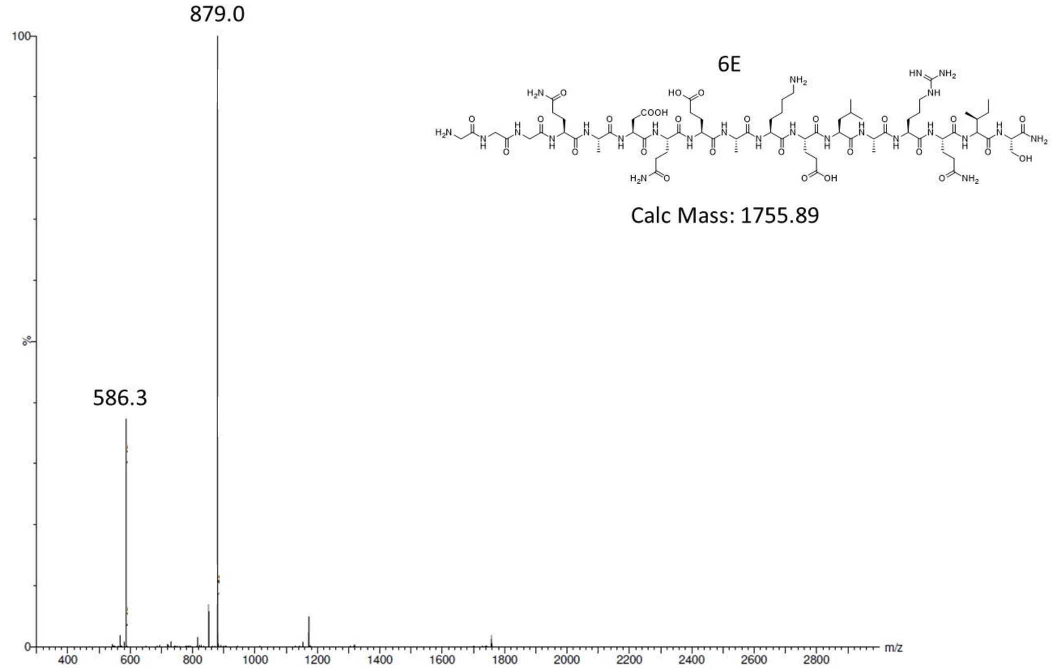


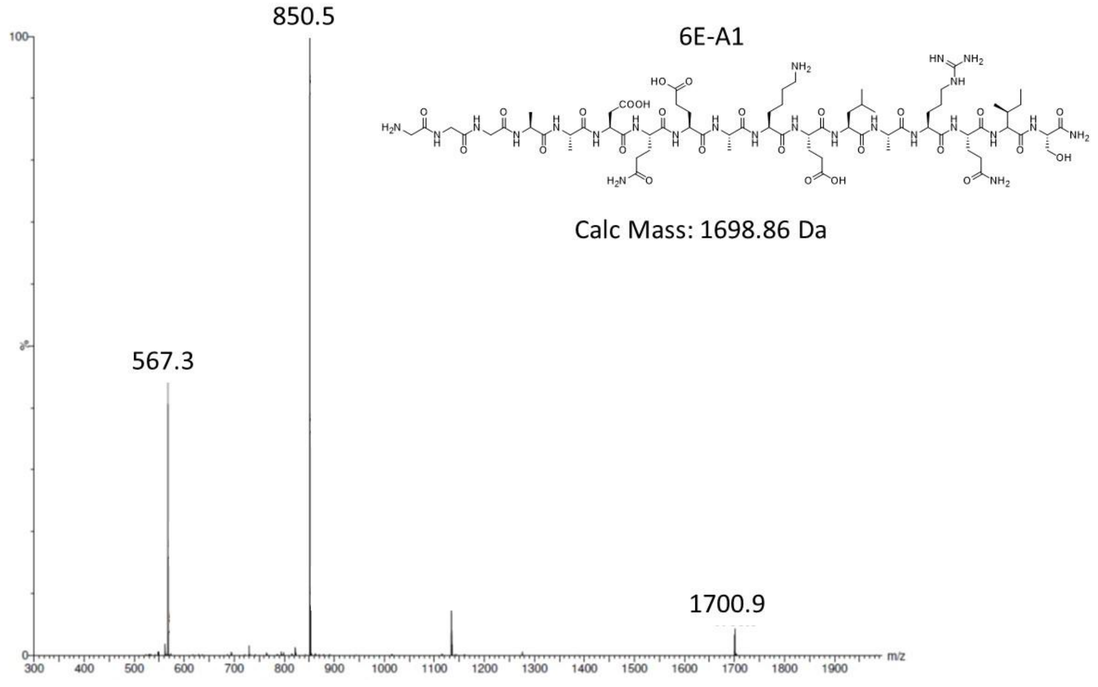

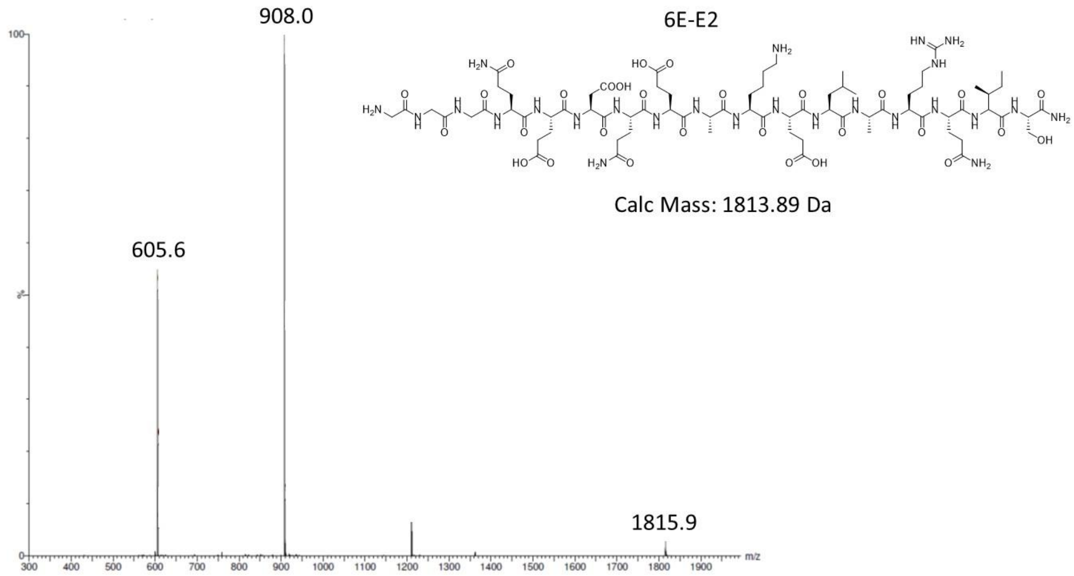
908.0
605.6
1815.9
6E-E2
Calc Mass: 1813.89 Da
m/z

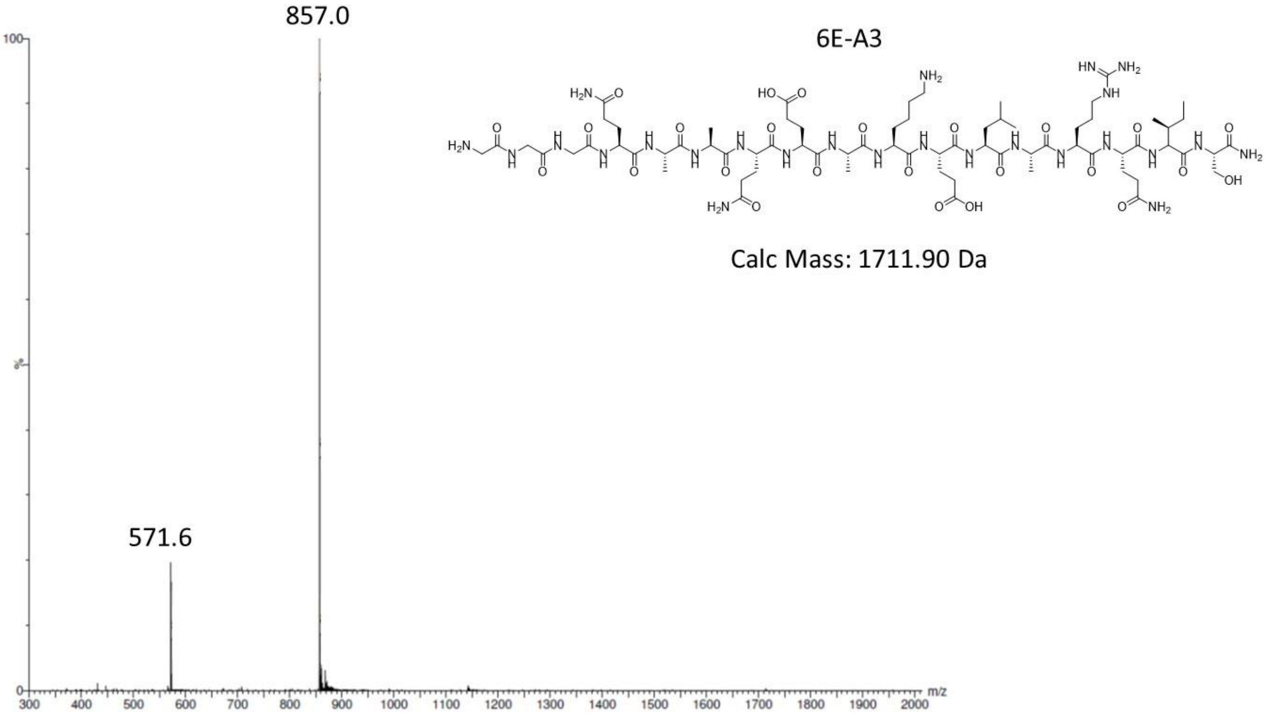
857.0
571.6
6E-A3
Calc Mass: 1711.90 Da
m/z

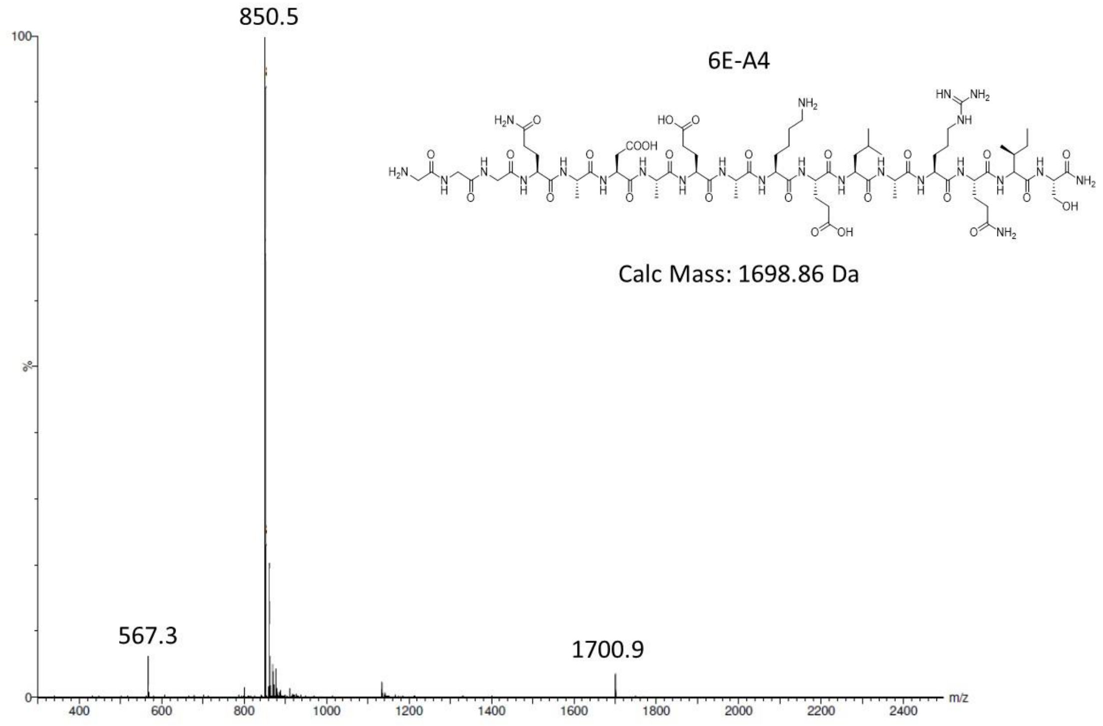
850.5
6E-A4
Calc Mass: 1698.86 Da
567.3
1700.9
100
%
0
m/z

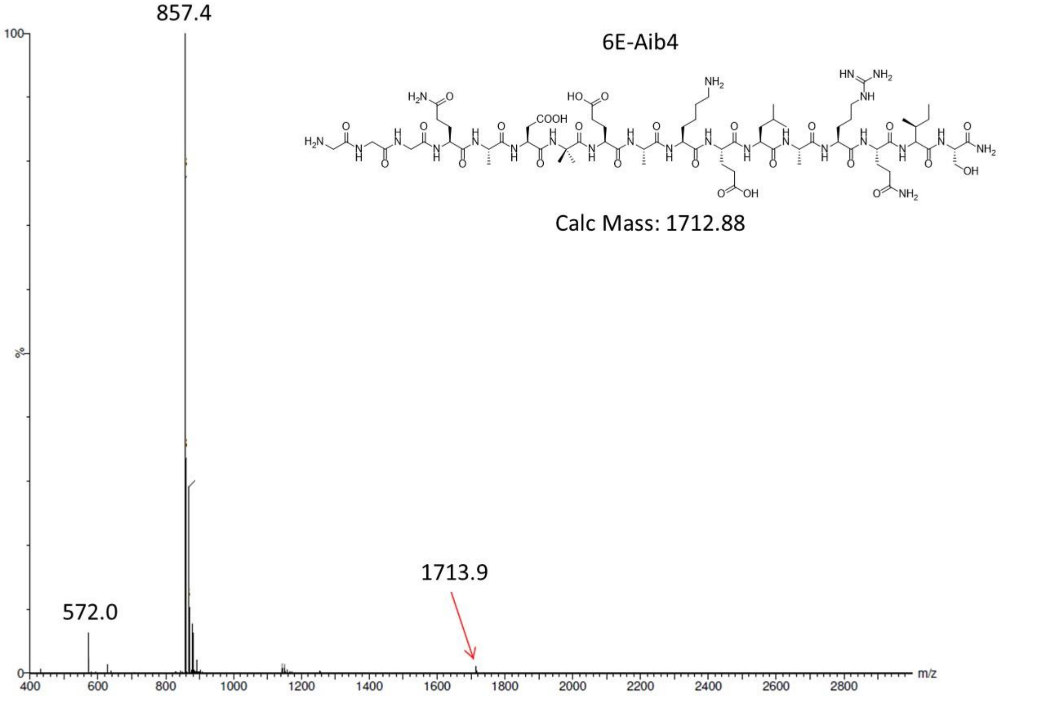
857.4
6E-Aib4
Calc Mass: 1712.88
572.0
1713.9
100
%
0
m/z

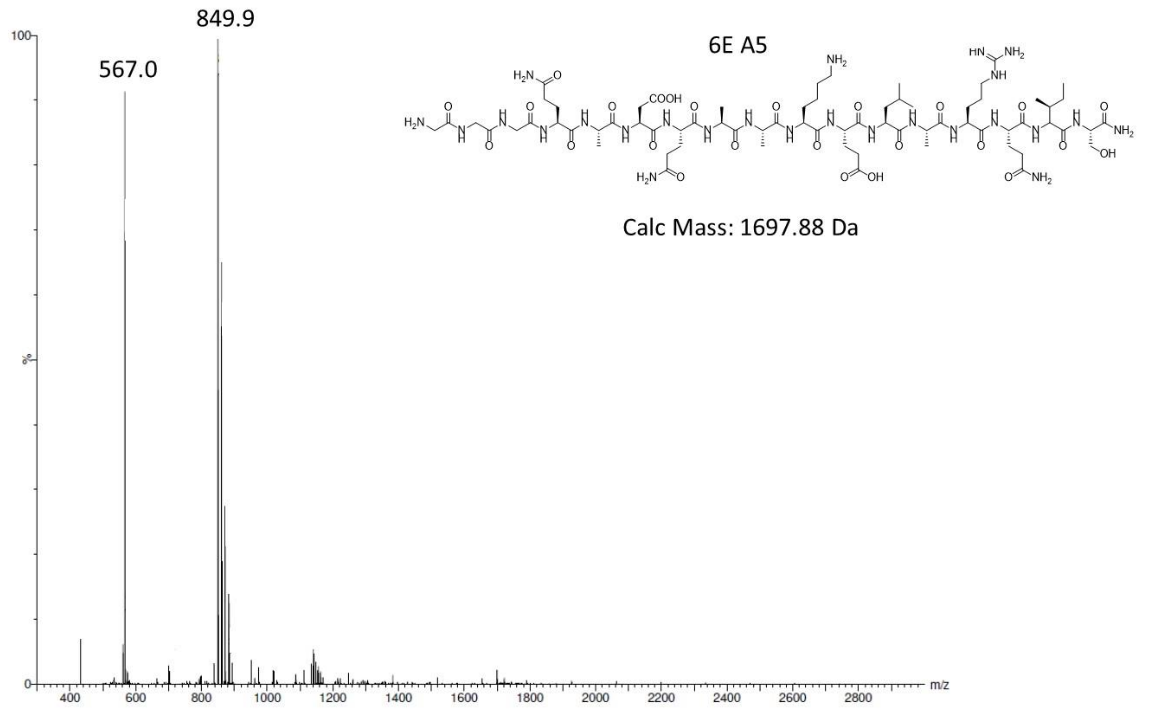

849.9
567.0
6E A5
Calc Mass: 1697.88 Da
100
%
0
m/z
400
600
800
1000
1200
1400
1600
1800
2000
2200
2400
2600
2800


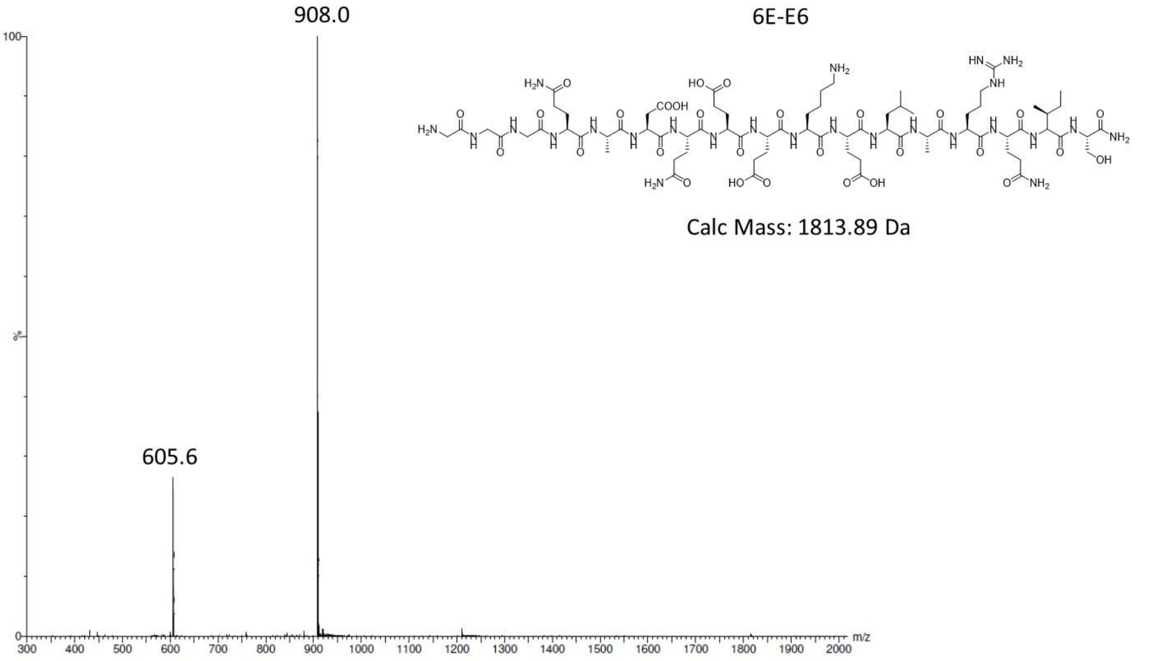

908.0
6E-E6
Calc Mass: 1813.89 Da
605.6
100
%
0
m/z
300
400
500
600
700
800
900
1000
1100
1200
1300
1400
1500
1600
1700
1800
1900
2000

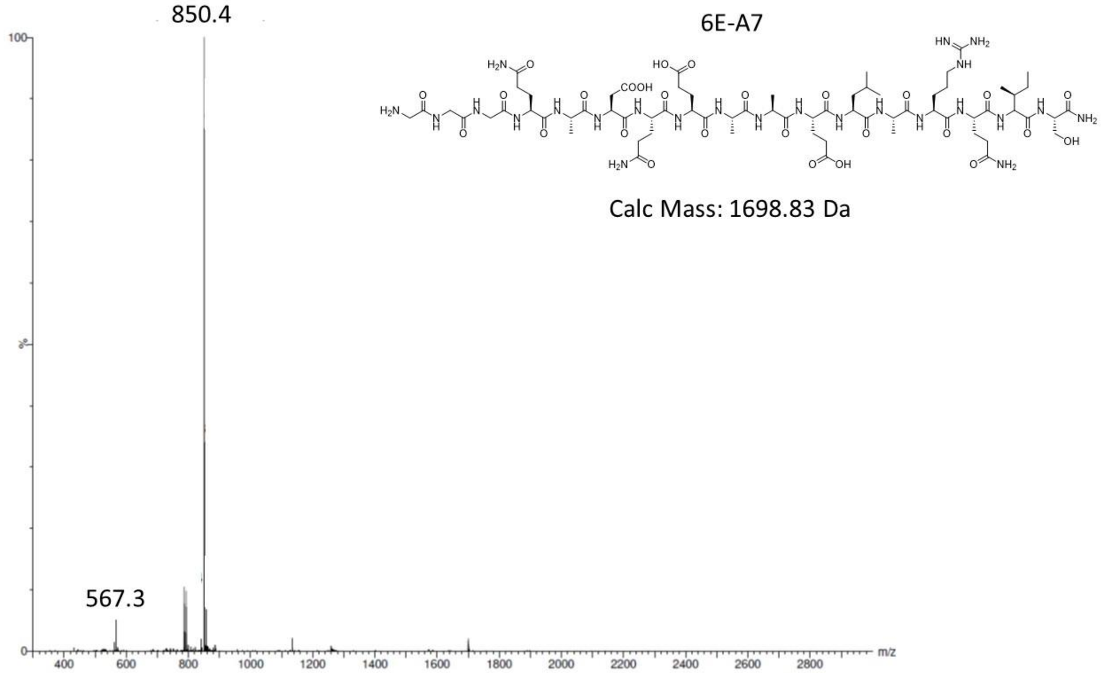
850.4
6E-A7
Calc Mass: 1698.83 Da
567.3
m/z

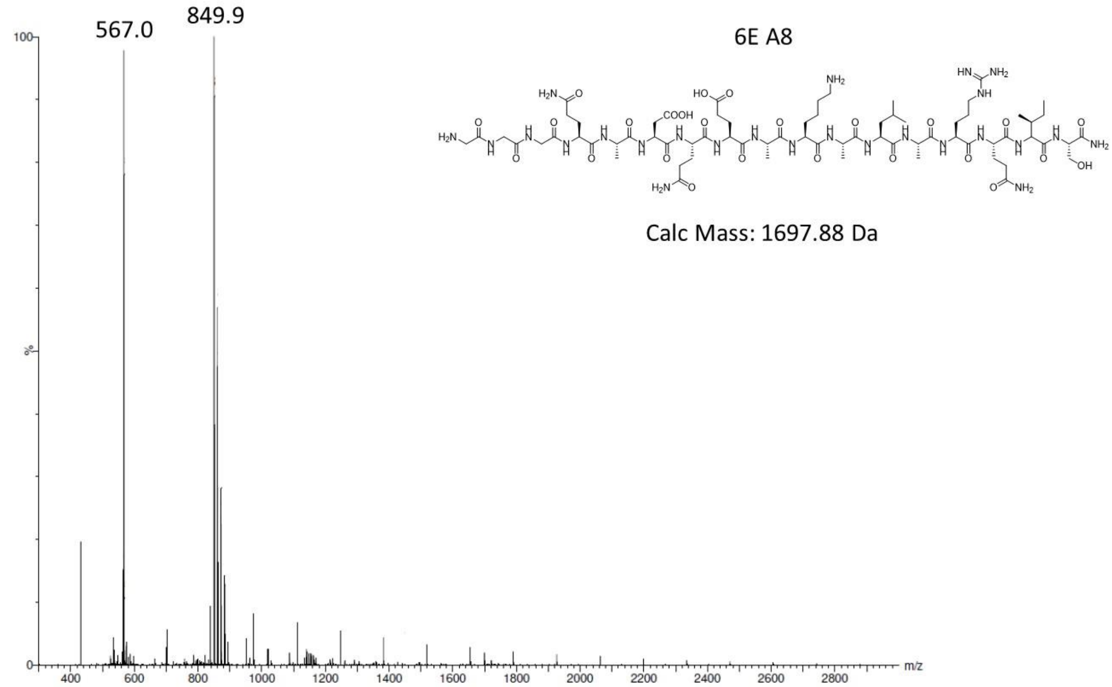
567.0
849.9
6E A8
Calc Mass: 1697.88 Da
m/z

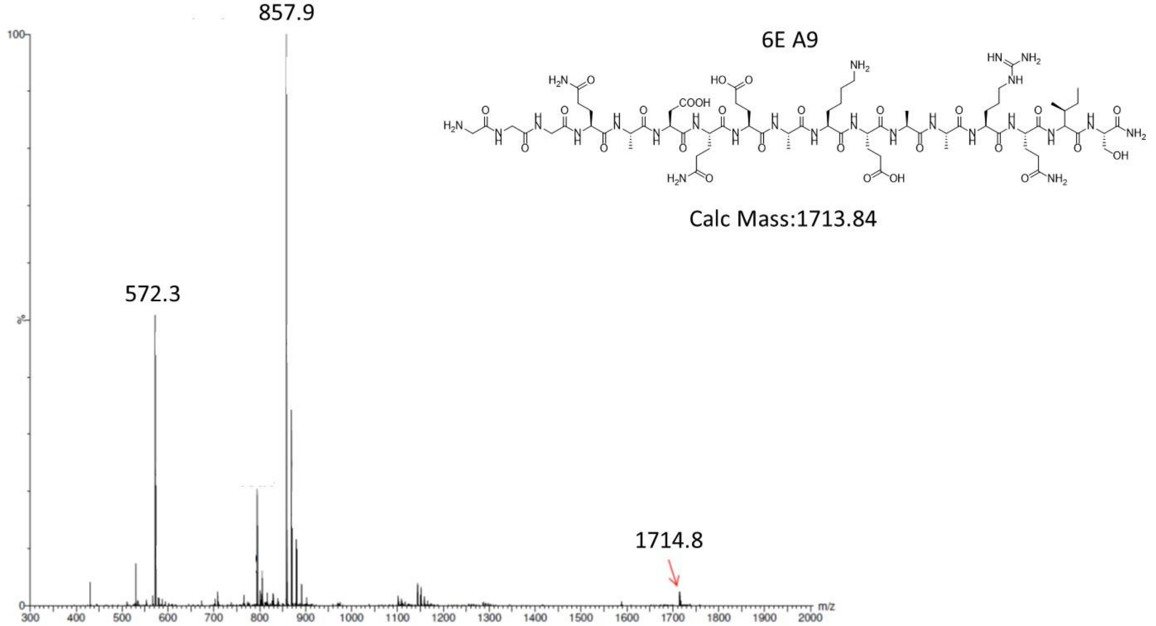


6E A9

Calc Mass:1713.84

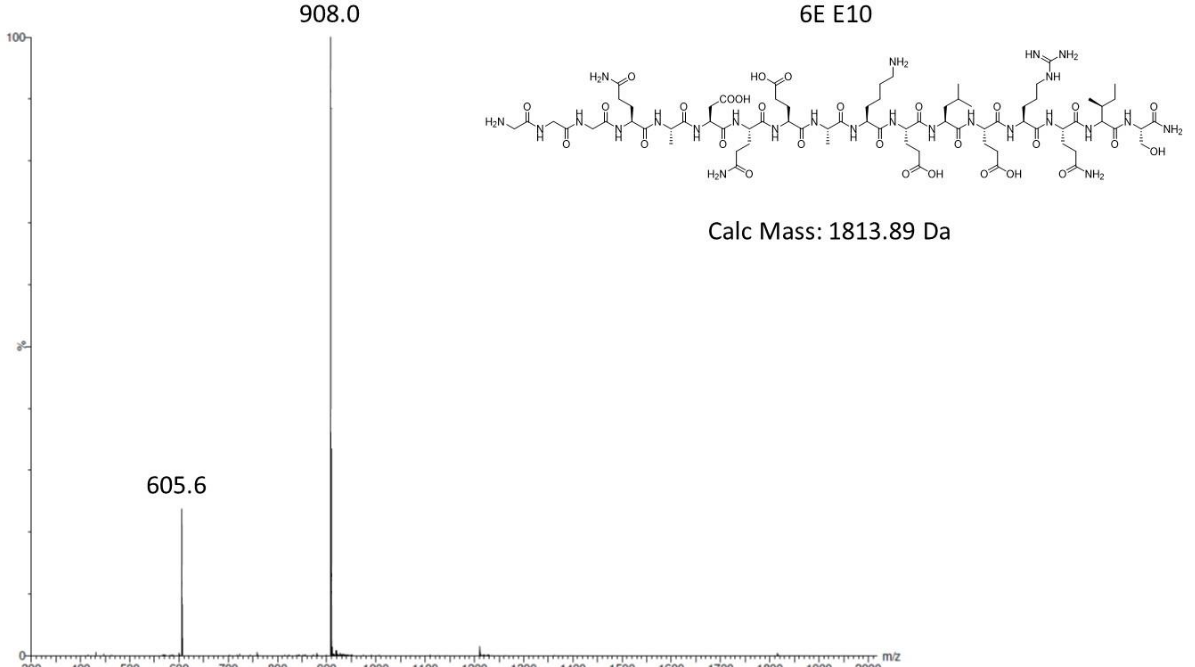


6E E10

Calc Mass: 1813.89 Da

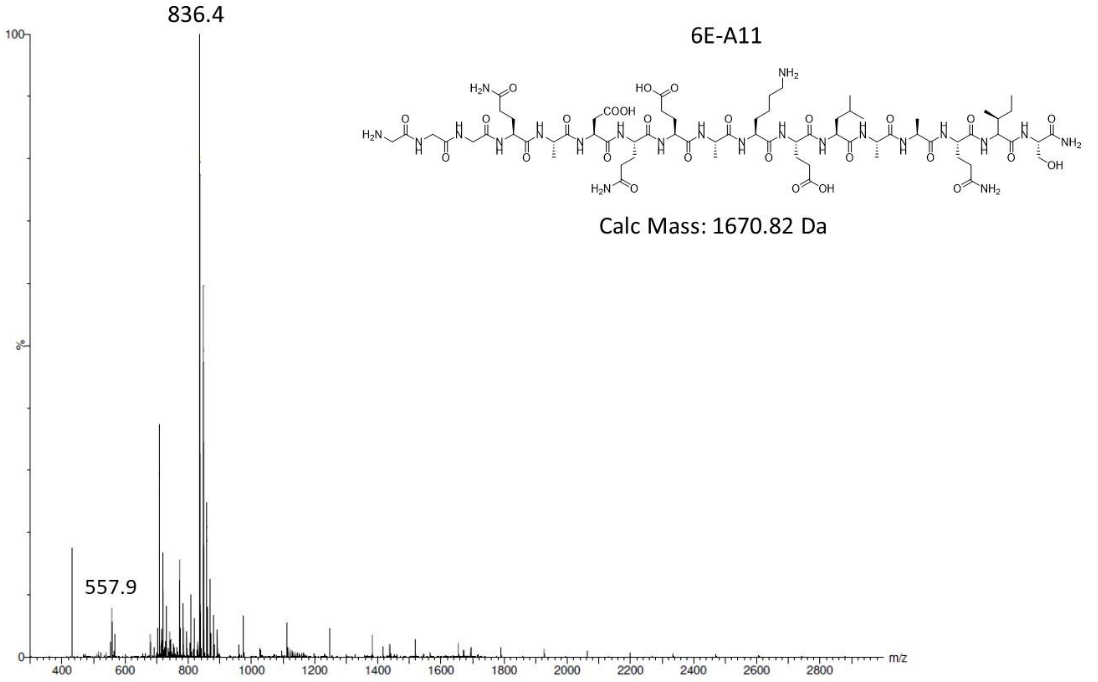

836.4
557.9
6E-A11
Calc Mass: 1670.82 Da
100
%
0
400 600 800 1000 1200 1400 1600 1800 2000 2200 2400 2600 2800
m/z


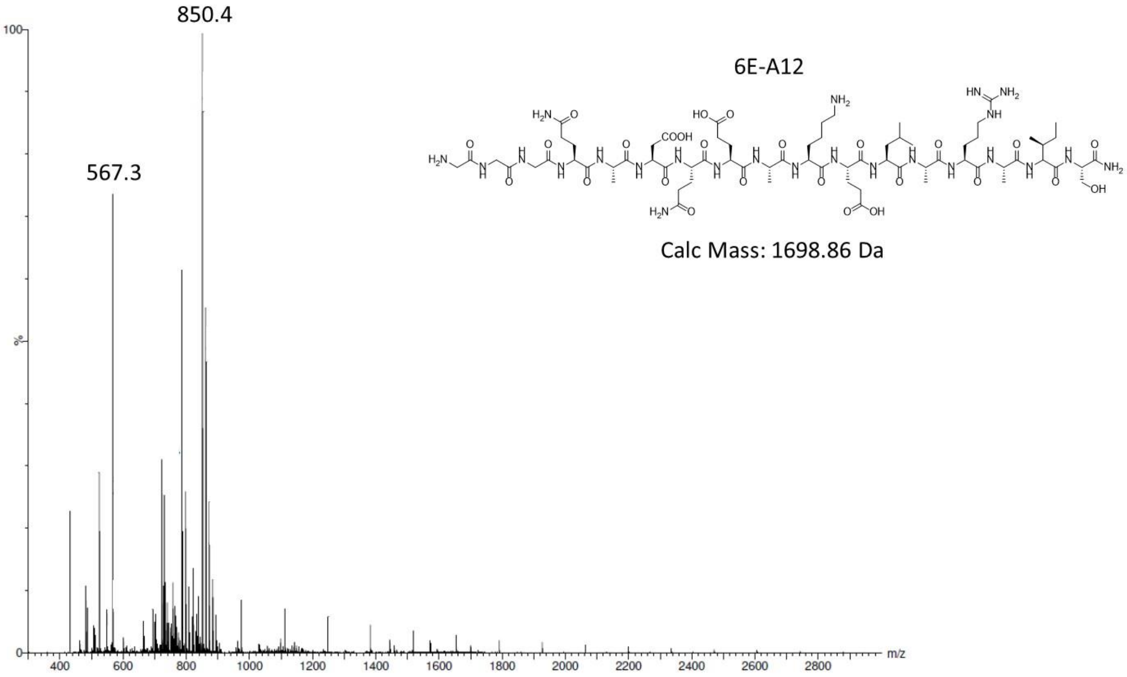

850.4
567.3
6E-A12
Calc Mass: 1698.86 Da
100
%
0
400 600 800 1000 1200 1400 1600 1800 2000 2200 2400 2600 2800
m/z

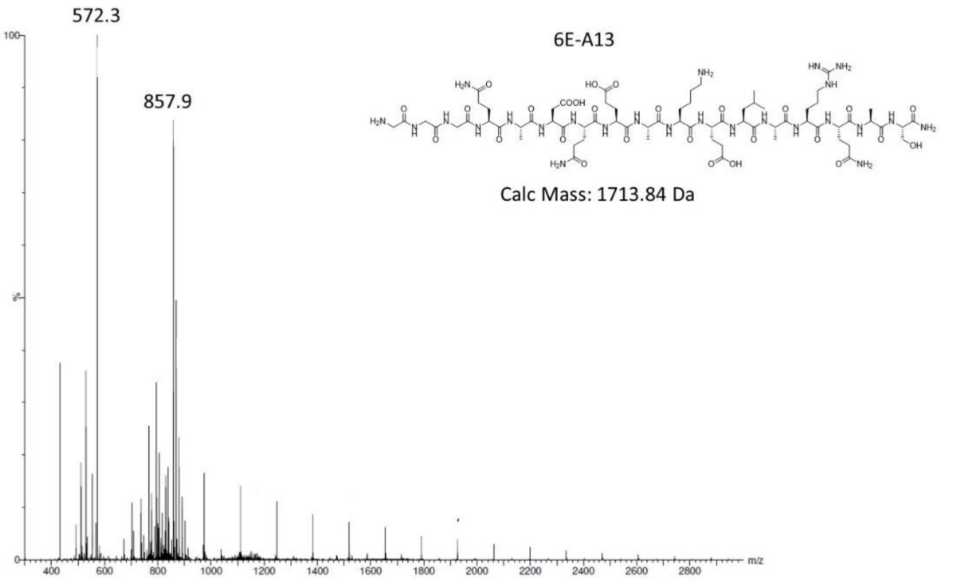
572.3
857.9
6E-A13
Calc Mass: 1713.84 Da
m/z

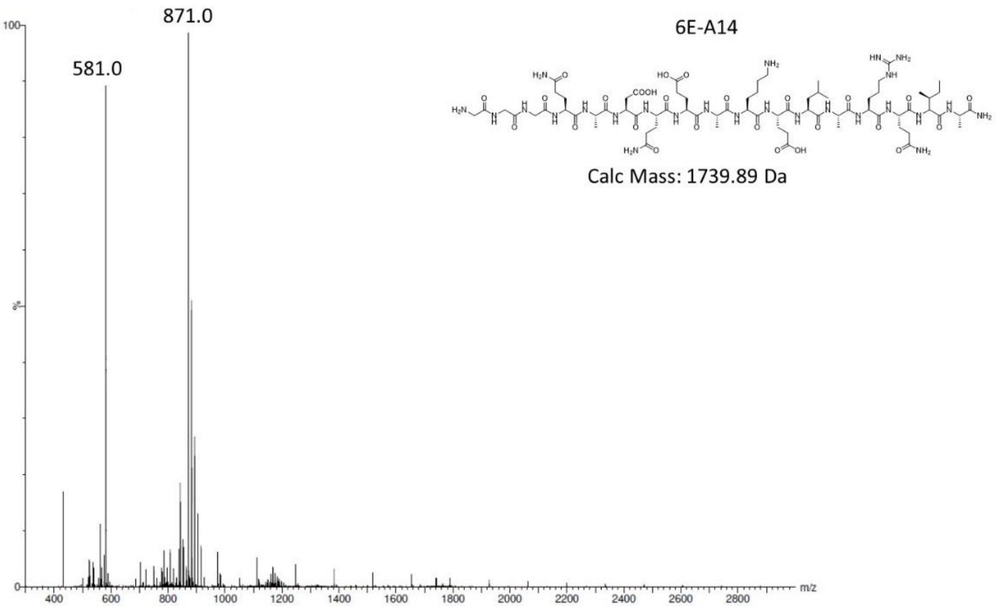
871.0
581.0
6E-A14
Calc Mass: 1739.89 Da
m/z

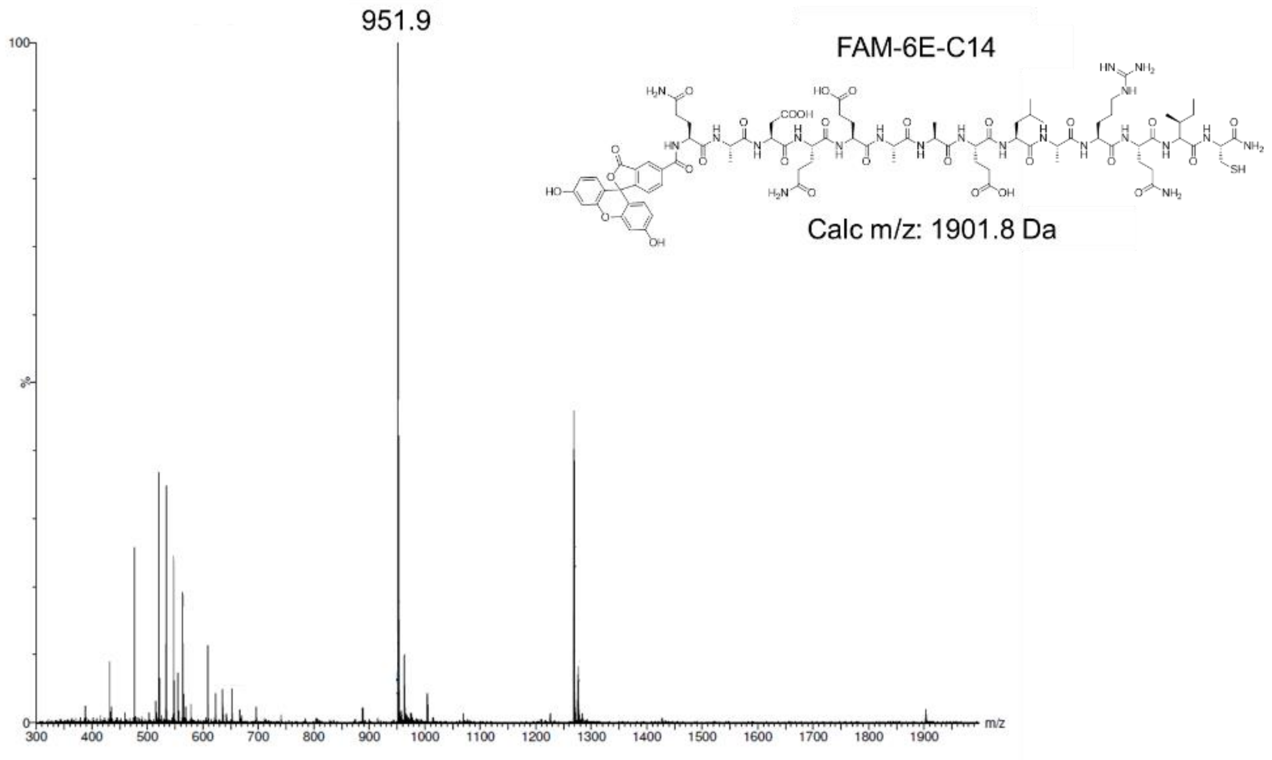
951.9
FAM-6E-C14
Calc m/z: 1901.8 Da
m/z
100
%
0
300
400
500
600
700
800
900
1000
1100
1200
1300
1400
1500
1600
1700
1800
1900

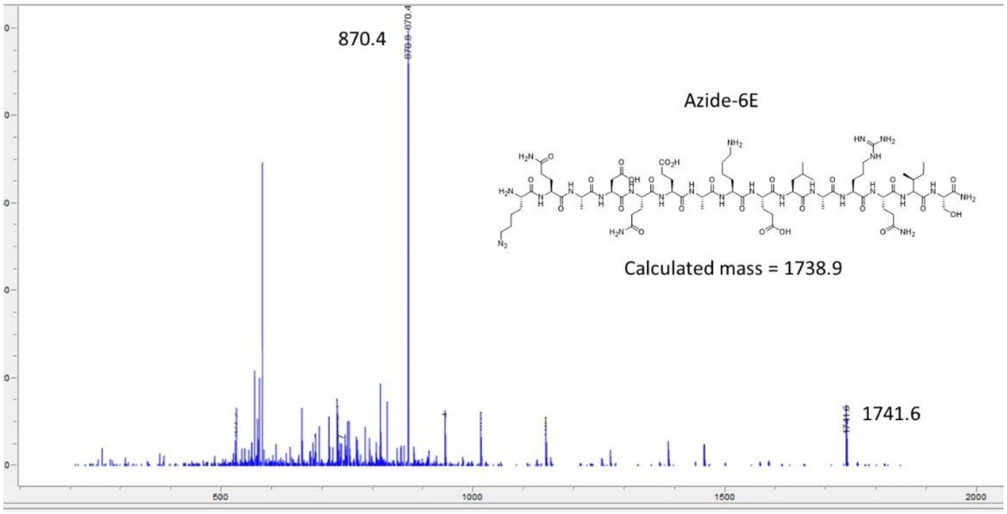
870.4
Azide-6E
Calculated mass = 1738.9
1741.6
500
1000
1500
2000

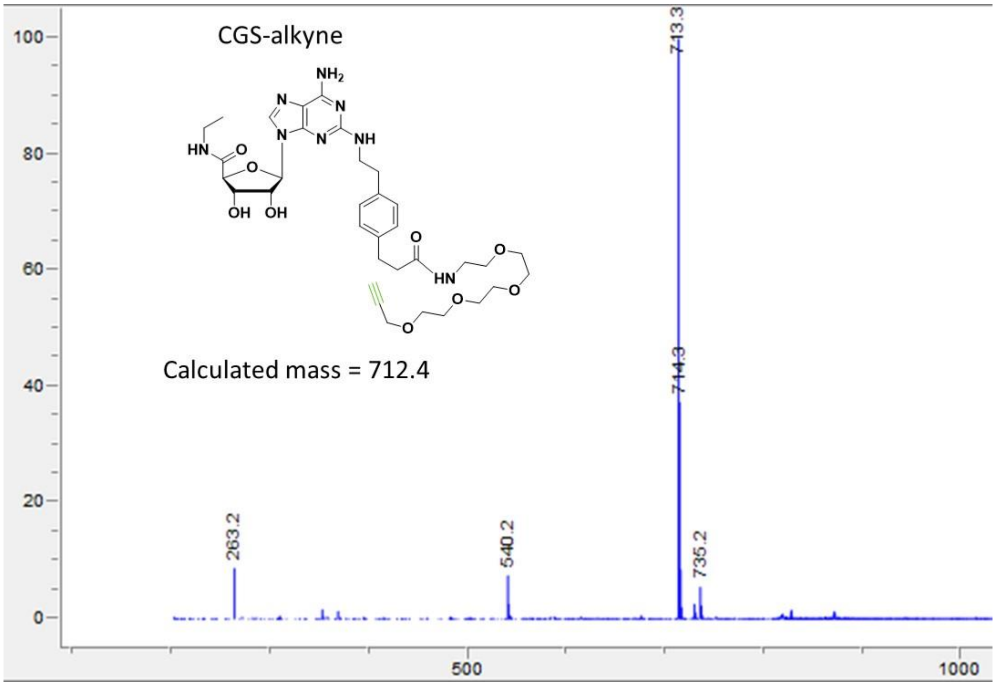
CGS-alkyne
Calculated mass = 712.4
713.3
714.3
263.2
540.2
735.2
500
1000

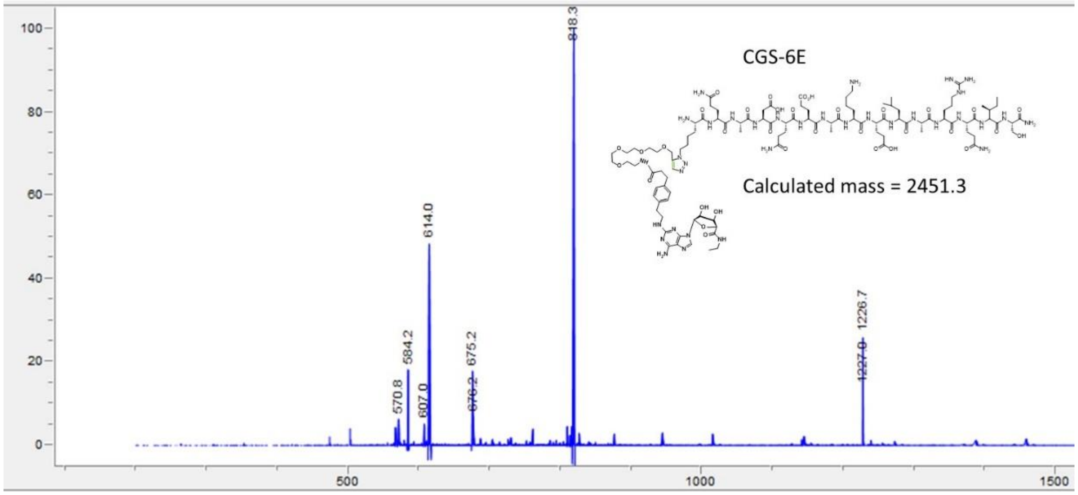
CGS-6E
Calculated mass = 2451.3
818.3
614.0
584.2
570.8
607.0
675.2
676.2
1226.7
1227.0
500
1000
1500

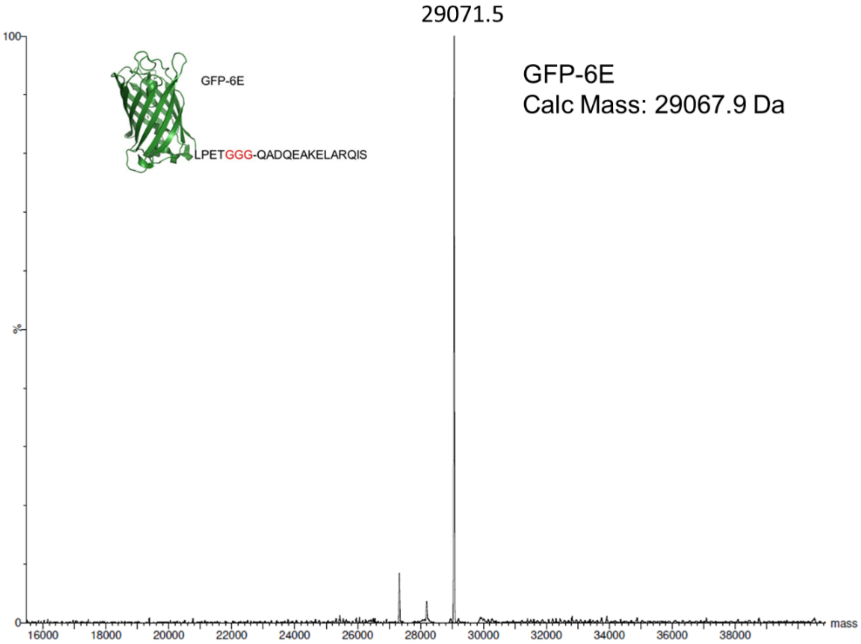


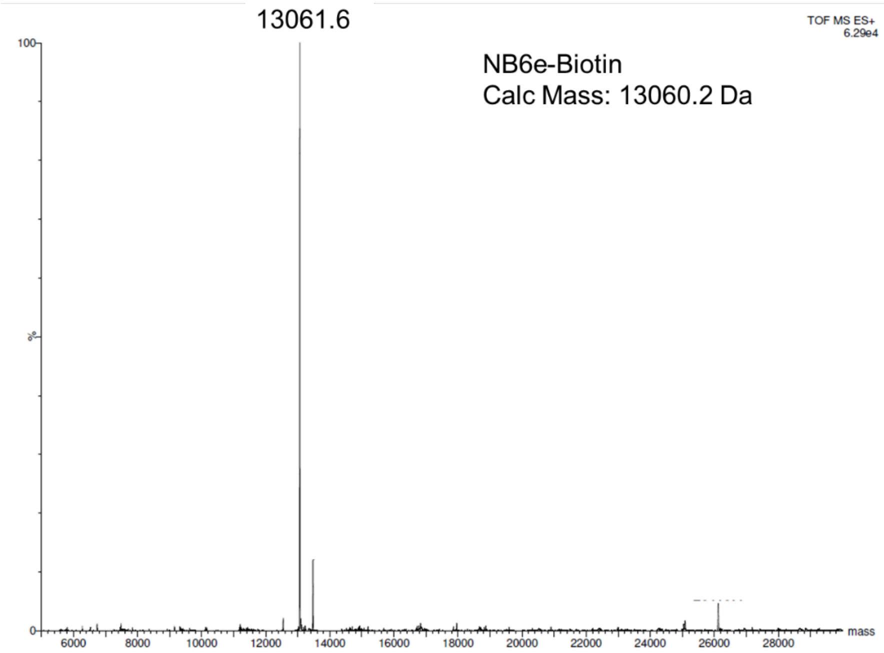


**Supporting Figure 1. Mass spectrometry characterization of compounds used in this study.** Compounds were analyzed by liquid chromatography/mass spectrometry as described in Methods.

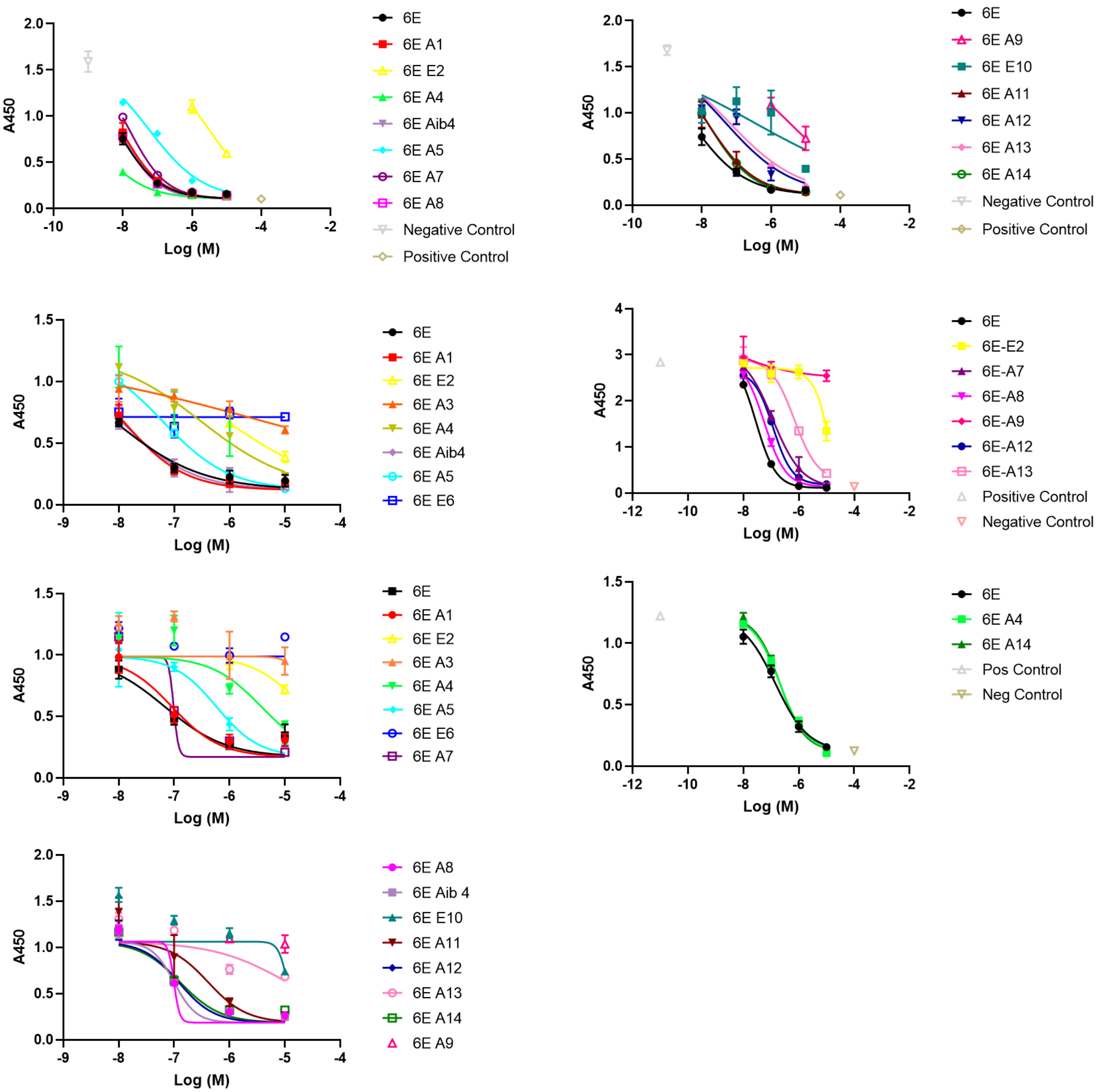


**Supporting Figure 2. Additional characterization of 6E peptide variants binding to $Nb_{6E}$.** Binding assays were performed using ELISA as described in Methods. Each graph corresponds to an independent experiment with data points corresponding to mean ± SD. Curves result from fitting a four-parameter sigmoidal dose-response model to the data (except for weak compounds, which show connected points).

| Peptide Analyte | $k_a$1 (1/M*s) | $k_d$1 (1/s) | $k_a$2 (1/M*s) | $k_d$2 (1/s) | $K_D$ (M) | Rmax top Dilution (RU) | Chi$^2$ |
|---|---|---|---|---|---|---|---|
| 6E Run 1 | 13220000 | 0.0668 | 0.02077 | 0.01109 | 1.759E-09 | 79.93 | 0.795 |
| 6E Run 2 | 3864000 | 0.1949 | 0.01048 | 0.01048 | 1.173E-08 | 40.57 | 0.514 |
| 6E Run 3 | 3915000 | 0.2201 | 0.0371 | 0.01123 | 1.306E-08 | 36.54 | 0.391 |
| 6E Run 4 | 5188000 | 0.08752 | 0.0292 | 0.00872 | 3.88E-09 | 95.33 | 2.29 |
| 6E Run 5 | 2948000 | 0.2559 | 0.02504 | 0.01448 | 3.18E-08 | 21.94 | 0.148 |

| Peptide Analyte | $k_a$1 (1/M*s) | $k_d$1 (1/s) | $k_a$2 (1/M*s) | $k_d$2 (1/s) | $K_D$ (M) | Rmax top Dilution (RU) | Chi$^2$ |
|---|---|---|---|---|---|---|---|
| 6E A5 Run 1 | 3549000 | 0.6254 | 0.02014 | 0.006126 | 4.109E-08 | 58.82 | 0.206 |
| 6E A5 Run 2 | 1919000 | 0.5573 | 0.01838 | 0.007947 | 8.764E-08 | 30.13 | 0.0734 |
| 6E A5 Run 3 | 4143000 | 0.533 | 0.01278 | 0.00358 | 2.814E-08 | 123.1 | 0.572 |

| Peptide Analyte | $k_a$1 (1/M*s) | $k_d$1 (1/s) | $k_a$2 (1/M*s) | $k_d$2 (1/s) | $K_D$ (M) | Rmax top Dilution (RU) | Chi$^2$ |
|---|---|---|---|---|---|---|---|
| 6E A9 Run 1 | 133400 | 0.7185 | 0.03654 | 0.001499 | 2.122E-07 | 32.84 | 0.0433 |
| 6E A9 Run 2 | 1005000 | 1.4132 | 0.004401 | 0.005049 | 7.613E-07 | 84.52 | 0.172 |

**Supporting Figure 3. Characterization of SPR analysis of 6E variant binding to Nb$_{6E}$.** Experiments were performed as described in Methods. Rmax top dilution refers to the maximum response (RU) of the most concentrated dilution in the resultant sensorgram. Chi$^2$ refers to the average of the square residuals, the difference between the experimental curves and the fit. A Chi$^2$ value less than 10% of the Rmax is a qualitative indicator of a good curve fit.

Independent replicate 2

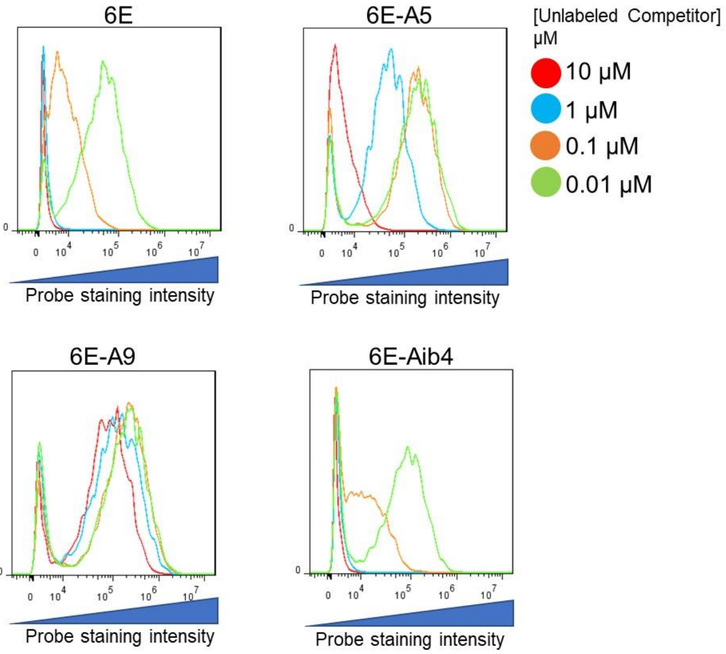


Independent replicate 3

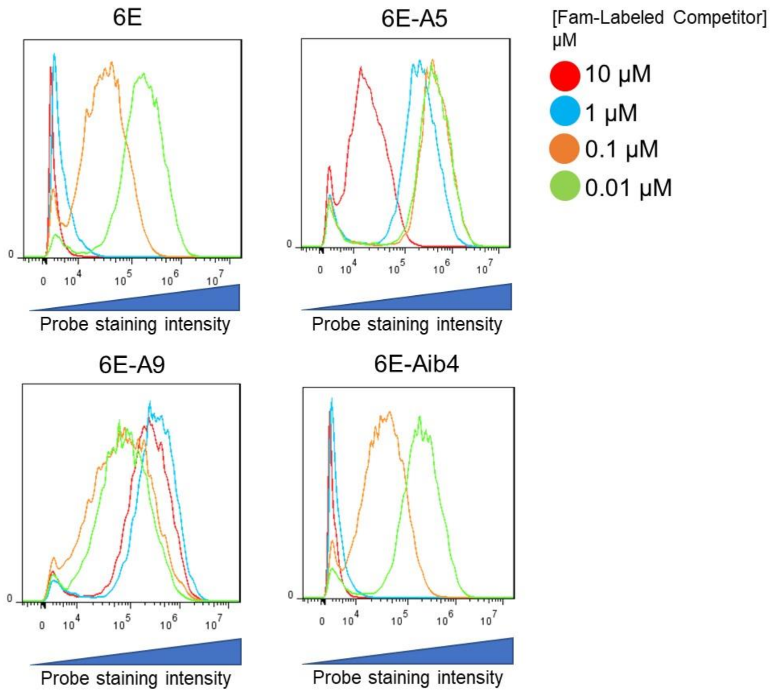


**Supporting Figure 4. Independent replicates for flow cytometry-based binding assays.** Assays were performed as described in Methods. Live cells were analyzed for the inhibition of tracer peptide (FAM-6E-C14) binding by named peptides at indicated concentrations. Histograms are shown with staining in the AF647 channel shown on the X-axis.

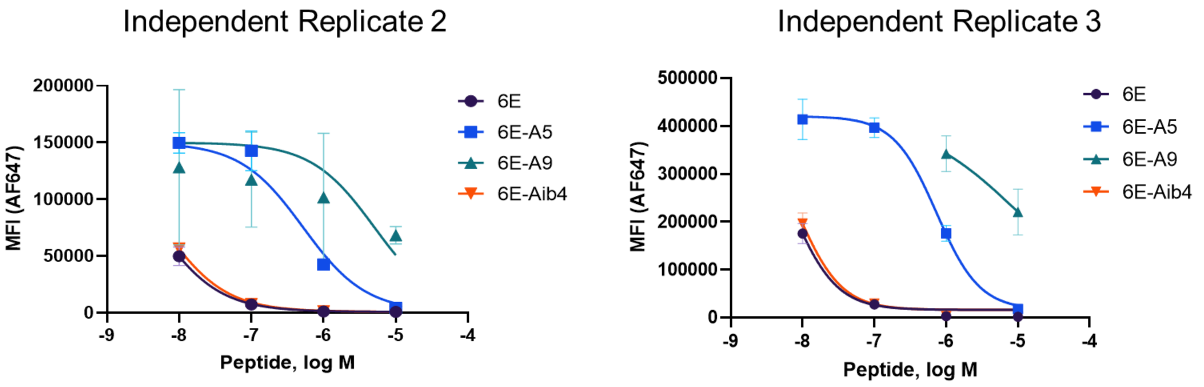


**Supporting Figure 5. Median fluorescence intensity (MFI) dose-response curves.** Data points represent the median fluorescence intensity derived from technical replicates of flow cytometry binding assays shown in Supporting Figure 4. Curves were fit to a 4-parameter sigmoidal dose-response model.

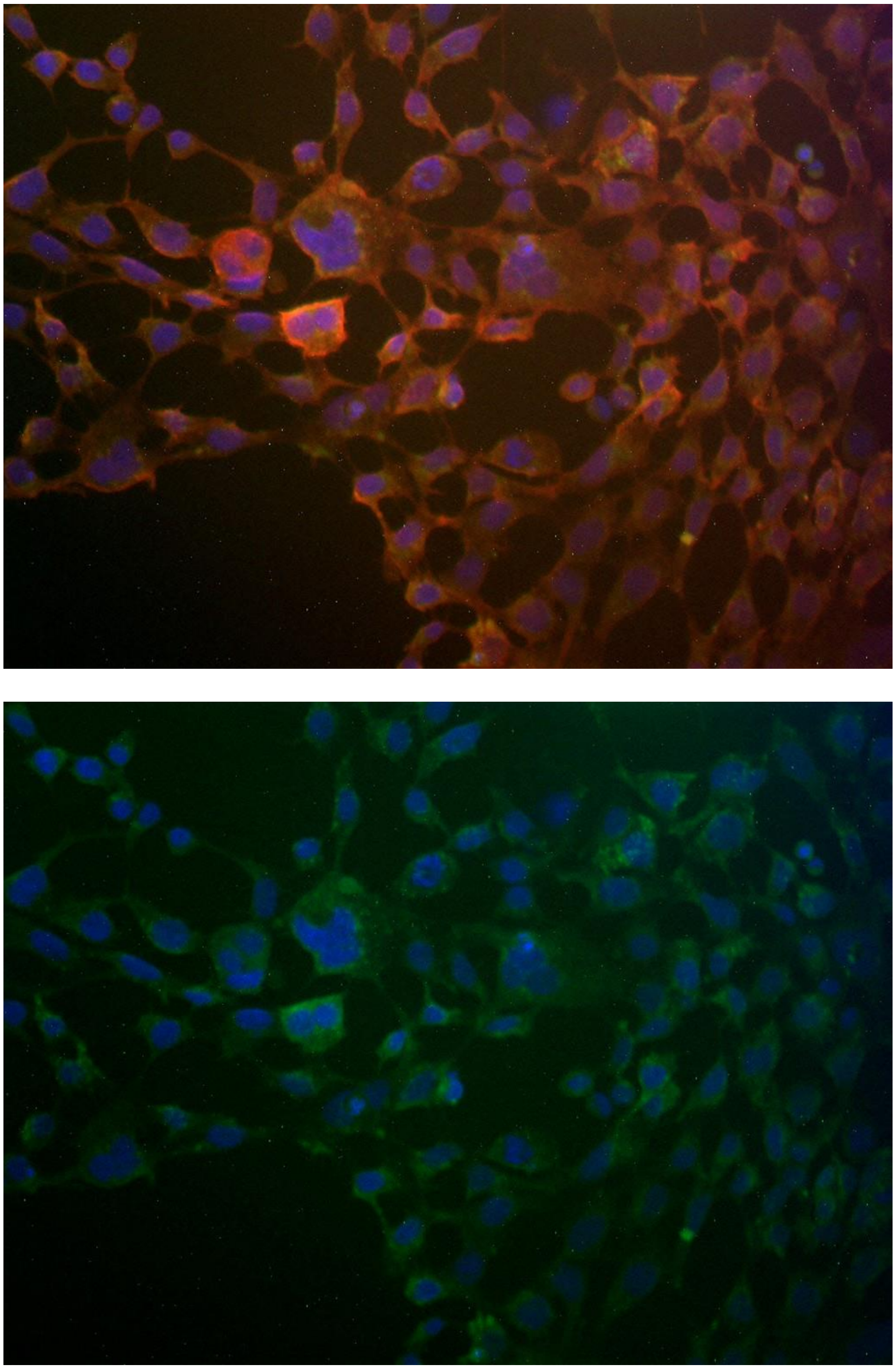

**Supporting Figure 6. Analysis of peptide labeling using fluorescence microscopy.** HEK293 cells expressing A2AR($Nb_{6E}$) were stained with FAM-6E-C14 (100 nM) and $Nb_{Alfa}$-TMR (300 nM). Note that this receptor contains an epitope tag in its extracellular portion bound by $Nb_{Alfa}$ (See supporting methods for sequence). Cells were washed, fixed, stained with DAPI and imaged at 20x magnification (See Supporting Methods). Top and bottom panels show an identical field of view with different fluorescence filter sets applied (top: FAM/TMR/DAPI; bottom: FAM/DAPI).

UBC6e (Modeled structure (non-6E peptide), 6E peptide, non-modeled portion of protein)

METRYNLKSPAVKRLMKEAAELKDPTDHYHAQPLEDNLFEWHFTVRGPPDSDFDGGVYHG
RIVLPPEYPMKPPSIILLTANGRFEVGKKICLSISGHHPETWQPSWSIRTALLAIIGFMP
TKGEGAIGSLDYTPEERRALAKKSQDFCCEGCGSAMKDVLLPLKSGSDSSQADQEAKELA
RQISFKAEVNSSGKTISESDLNHSFSLTDLQDDIPTTFQGATASTSYGLQNSSAASFHQP
TQPVAKNTSMSPRQRRAQQQSQRRLSTSPDVIQGHQPRDNHTDHGGSAVLIVILTLALAA
LIFRRIYLANEYIFDFEL

$Nb_{6E}$ (CDR1/CDR2/CDR3)-linker-6E peptide (linker not shown in structure)

QVQLQESGGGLVQPGGSLRLSCAASGFVFENSAMAWYRQAPGKERELIAVIGTTFIKLAESVK
GRFTISRDNAKSTVYLQMNNLKPEDTAVYYCSKSGAYWGQGTQVTVSSGGGSGGGSGGGSG
GGSGGGSGGGSGGGSGGGSGGGSQADQEAKELARQIS

**Supporting Figure 7. Annotated sequences for structural models.** Structural models presented in the main text were generated through input of these sequences into Alphafold2 Colab notebook. Structures were visualized using Pymol and colored according to the annotation above.

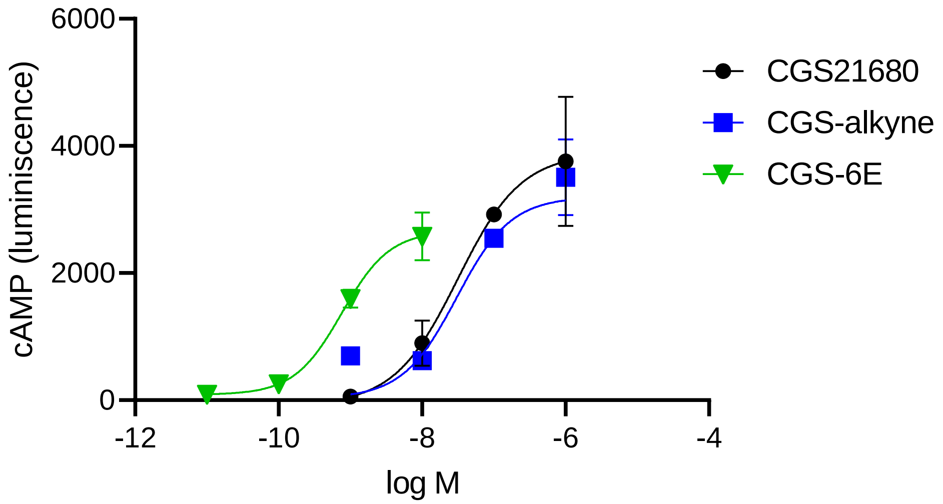

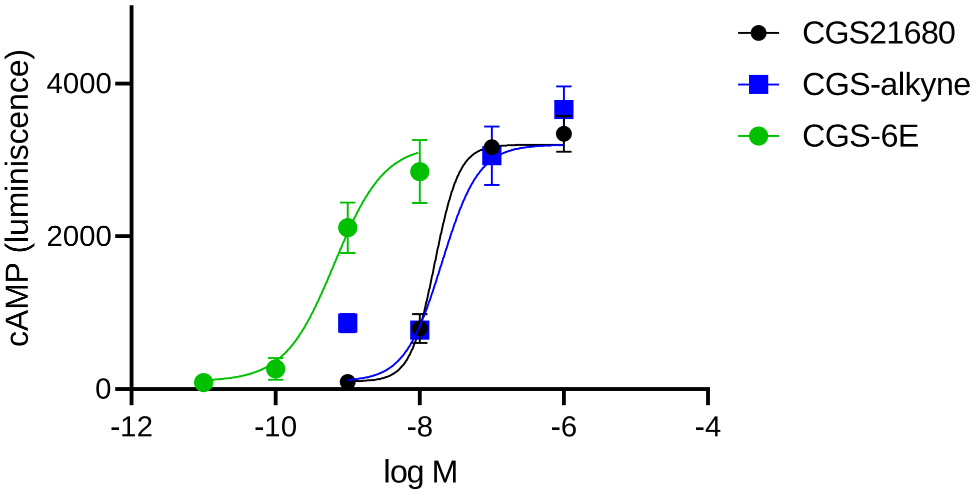


**Supporting Figure 8. Replicate data for activation of A2AR($Nb_{6E}$) signaling in HEK293 cells.** Data sets correspond to dose-response curves cAMP responses generated upon addition of ligands to cells for 12 minutes. Curves were generated by fitting data to a 4-parameter sigmoidal dose-response. Each graph is an independent experiment (separate from that shown in Figure 6B). Data points correspond to mean ± SD from technical replicates within a single experiment.

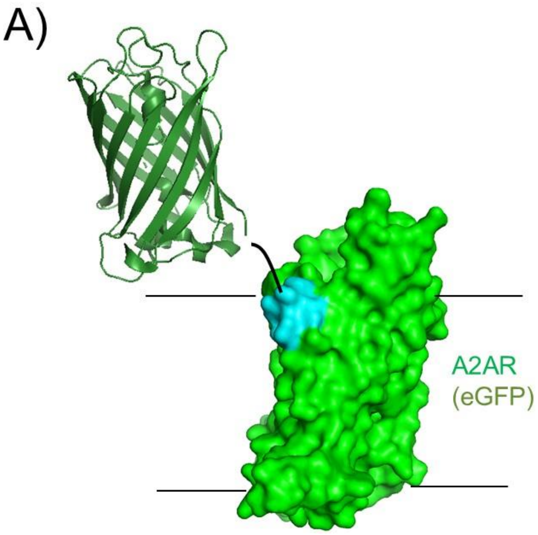


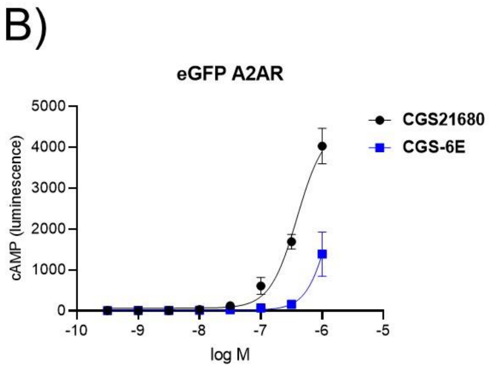


| | $EC_{50}$, nM | SD | Fold |
|---|---|---|---|
| CGS21680 | 420 | 130 | 1.0 |
| CGS-6E | 3200 | 1700 | 7.6 |

**Supporting Figure 9. Evaluation of pharmacological properties of CGS-6E on an A2AR(eGFP) fusion protein.** A) Schematic depiction of the receptor construct used for experiments in this figure. B) (Top) Representative dose-response curve for the action of indicated compounds for inducing cAMP responses on cells expressing A2AR(eGFP). Data points represent mean ± SD from technical replicates in a single experiment. (Bottom) Tabulation of compound agonist potency parameters. Indicated parameters represent mean ± SD from three independent replicate experiments. “Fold” indicates the $EC_{50}$ value normalized to that of CGS21680.

**Supporting Methods.**

**Data analysis.** Average values and uncertainties correspond to mean ± standard deviation. Dose-response data were fit to 4-parameter sigmoidal dose-response model.

**Cell culture and cell lines.** HEK293 cells (ATCC CRL-1573) were transfected with a cAMP responsive luciferase variant[1] and the receptor of interest to provide clonal cell lines that stably express cAMP biosensor and receptor and can be grown without selection antibiotic. All cell lines were cultured in DMEM supplemented with 10% fetal bovine serum and penicillin/streptomycin. Cells were checked for mycoplasma contamination.

**Measurement of receptor activation using cAMP-responsive luciferase expression.** Assays were run according to previously described protocols[2]. Briefly, cells were plated in a white walled, clear bottom 96 well plate (Corning #3610) and grown to confluency. Growth medium was removed and $CO_2$ independent medium containing luciferin (0.5 mM) was added until a stable background reading was obtained (~10 minutes). 10x stocks of ligands were then added to wells and cAMP (luminescence) responses were measured for every 2 minutes for 12 minutes after ligand addition (Biotek Neo2 plate reader). The response recorded 12 minutes after ligand addition was used for construction of dose-response curves.

**Protein expression plasmids.** Nanobodies were expressed in *E. coli* from pET26b(+) plasmids encoding a sequence corresponding to pelB leader-nanobody-LPETGG-His6 under the control of Lac repressor with ampicillin resistance.

**Receptor Plasmids.** Custom plasmids were ordered from VectorBuilder. Plasmids included elements encoding G418 resistance and a CAG promoter for the protein of interest. Insert sequences are listed below here.

HGH secretion tag-alfa tag-$Nb_{6E}$-human A2AR

Amino acid

MATGSRTSLLLAFGLLCLPWLQEGSAFPTIPLSGSEPSRLEEELRRRLTEPSGSMAQVQLQES
GGGLVQPGGSLRLSCAASGFVFENSAMAWYRQAPGKERELIAVIGTTFIKLAESVKGRFTISRD
NAKSTVYLQMNNLKPEDTAVYYCSKSGAYWGQGTQVTVSSGGLPETGGSGGMPIMGSSVYIT
VELAIAVLAILGNVLVCWAVWLNSNLQNVTNYFVVSLAAADIAVGVLAIPFAITISTGFCAACHGC
LFIACFVLVLTQSSIFSLLAIAIDRYIAIRIPLRYNGLVTGTRAKGIIAICWVLSFAIGLTPMLGWNNC
GQPKEGKNHSQGCGEGQVACLFEDVVPMNYMVYFNFFACVLVPLLLMLGVYLRIFLAARRQL
KQMESQPLPGERARSTLQKEVHAAKSLAIIVGLFALCWLPLHIINCFTFFCPDCSHAPLWLMYLA
IVLSHTNSVVNPFIYAYRIREFRQTFRKIIRSHVLRQQEPFKAAGTSARVLAAHGSDGEQVSLRL
NGHPPGVWANGSAPHPERRPNGYALGLVSGGSAQESQGNTGLPDVELLSHELKGVCPEPPG
LDDPLAQDGAGVS*

Nucleotide

atggcaacaggatcaaggacatccttgcttctcgcattcggccttctctgcctgccttggctgcaagagggtagcgcatttcctaccatac
ccttgtccggaagtgaaccctctaggctggaggaggaattgagacgccggttgacagagccctccggatccatggcacaggtccag
cttcaagagtccggtggtggtctggtgcagcccggcggttcactgcgccttagctgtgcggcaagcggctttgtttttgaaaacagcgca
atggcgtggtaccggcaggccccggggaaagagcgggagctcattgctgtgattggcactacttttattaaacttgcagaaagcgtta
agggccggtttacaatcagcagggataacgcgaagagcacagtatatctgcaaatgaacaacttgaagcctgaggacactgcagt

ctactattgtagtaagtctggtgcgtattggggacagggtacacaggtgaccgtcagctctggtggtcttcccgaaaccgggggttccgg
gggaatgccaatcatggggagctctgtatatatcacggtcgaactcgccattgctgtactggctattttgggcaacgttttggtttgctggg
cggtatggcttaatagcaatcttcaaaatgtgactaattacttcgtggtctcccttgctgcagccgatatcgccgttggtgttttggcgatacc
attcgcgatcaccatctctaccggcttttgtgctgcctgccatggctgcttgttcattgcttgttttgttttggtgctcacgcaatccagtatcttca
gtctcctcgcaatcgctatagatagatatatagcgatacgaatcccattgaggtacaacggactcgtaacaggaacaagagcgaaa
ggcataatagcaatttgctgggtgctctccttcgctattggactgaccccaatgctgggatggaacaattgtgggcaaccaaaagagg
gaaaaaaccacagccagggttgcggagagggtcaagtcgcttgcctttttgaagacgttgtcccaatgaattacatggtatattttaatttt
tttgcgtgcgtattggtacctttgctgctcatgctcggagtctatcttagaatatttcttgcggctcgacgacaactgaagcaaatggagtca
cagcctttgcctggggagagggcaagatctacccttcagaaggaggtgcatgcagcaaaaagccttgccataatcgtaggcctgttc
gctctttgctggttgccacttcatatcatcaactgtttcacgttcttttgtccagattgcagtcatgcgccgttgtggttgatgtatctggcaatcg
tgctgtcacataccaattcagttgttaacccatttatctatgcctatcgcatccgggagtttcgccaaacttttaggaagataatcaggagtc
atgtcctgcggcagcaggaaccatttaaagcggcagggacatccgcacgcgttttggcggcgcatggatcagacggggagcaagt
atcattgcgcctcaacgggcacccccctggtgtttgggctaatggatcagccccgcacccagaacggcggcctaatggttacgccttg
ggcttggtgagcgggggctccgcccaggagtcacaggggaacactggccttcctgacgtggaactcttgagtcacgagctgaaggg
cgtttgcccagagcctcccggactcgatgatccactggctcaagatggggctggcgtgtcatga

Amino acid

HGH secretion tag-alfa tag-eGFP-human A2AR

MATGSRTSLLLAFGLLCLPWLQEGSAFPTIPLSGSEPSRLEEELRRRLTEPSGSMVSKGEELFT
GVVPILVELDGDVNGHKFSVSGEGEGDATYGKLTLKFICTTGKLPVPWPTLVTTLTYGVQCFSR
YPDHMKQHDFFKSAMPEGYVQERTIFFKDDGNYKTRAEVKFEGDTLVNRIELKGIDFKEDGNIL
GHKLEYNYNSHNVYIMADKQKNGIKVNFKIRHNIEDGSVQLADHYQQNTPIGDGPVLLPDNHYL
STQSALSKDPNEKRDHMVLLEFVTAAGITLGMDELYKGSGGMPIMGSSVYITVELAIAVLAILGN
VLVCWAVWLNSNLQNVTNYFVVSLAAADIAVGVLAIPFAITISTGFCAACHGCLFIACFVLVLTQS
SIFSLLAIAIDRYIAIRIPLRYNGLVTGTRAKGIIAICWVLSFAIGLTPMLGWNNCGQPKEGKNHSQ
GCGEGQVACLFEDVVPMNYMVYFNFFACVLVPLLLMLGVYLRIFLAARRQLKQMESQPLPGER
ARSTLQKEVHAAKSLAIIVGLFALCWLPLHIINCFTFFCPDCSHAPLWLMYLAIVLSHTNSVVNPFI
YAYRIREFRQTFRKIIRSHVLRQQEPFKAAGTSARVLAAHGSDGEQVSLRLNGHPPGVWANGS
APHPERRPNGYALGLVSGGSAQESQGNTGLPDVELLSHELKGVCPEPPGLDDPLAQDGAGVS
*

Nucleotide

atggcaacaggatcaaggacatccttgcttctcgcattcggccttctctgcctgccttggctgcaagagggtagcgcatttcctaccatac
ccttgtccggaagtgaaccctctaggctggaggaggaattgagacgccggttgacagagccctccggatccatggtgagcaagggc
gaggagctgttcaccggggtggtgcccatcctggtcgagctggacggcgacgtaaacggccacaagttcagcgtgtccggcgagg
gcgagggcgatgccacctacggcaagctgaccctgaagttcatctgcaccaccggcaagctgcccgtgccctggcccaccctcgtg
accaccctgacctacggcgtgcagtgcttcagccgctaccccgaccacatgaagcagcacgacttcttcaagtccgccatgcccgaa
ggctacgtccaggagcgcaccatcttcttcaaggacgacggcaactacaagacccgcgccgaggtgaagttcgagggcgacacc
ctggtgaaccgcatcgagctgaagggcatcgacttcaaggaggacggcaacatcctggggcacaagctggagtacaactacaac
agccacaacgtctatatcatggccgacaagcagaagaacggcatcaaggtgaacttcaagatccgccacaacatcgaggacggc
agcgtgcagctcgccgaccactaccagcagaacacccccatcggcgacggccccgtgctgctgcccgacaaccactacctgagc
acccagtccgccctgagcaaagaccccaacgagaagcgcgatcacatggtcctgctggagttcgtgaccgccgccgggatcactct
cggcatggacgagctgtacaagggttccgggggaatgccaatcatggggagctctgtatatatcacggtcgaactcgccattgctgta
ctggctattttgggcaacgttttggtttgctgggcggtatggcttaatagcaatcttcaaaatgtgactaattacttcgtggtctcccttgctgc
agccgatatcgccgttggtgttttggcgataccattcgcgatcaccatctctaccggcttttgtgctgcctgccatggctgcttgttcattgctt
gttttgttttggtgctcacgcaatccagtatcttcagtctcctcgcaatcgctatagatagatatatagcgatacgaatcccattgaggtaca

acggactcgtaacaggaacaagagcgaaaggcataatagcaatttgctgggtgctctccttcgctattggactgaccccaatgctggg
atggaacaattgtgggcaaccaaaagagggaaaaaaccacagccagggttgcggagagggtcaagtcgcttgcctttttgaagac
gttgtcccaatgaattacatggtatattttaattttttgcgtgcgtattggtacctttgctgctcatgctcggagtctatcttagaatatttcttgcg
gctcgacgacaactgaagcaaatggagtcacagcctttgcctggggagagggcaagatctacccttcagaaggaggtgcatgcag
caaaaagccttgccataatcgtaggcctgttcgctctttgctggttgccacttcatatcatcaactgtttcacgttcttttgtccagattgcagtc
atgcgccgttgtggttgatgtatctggcaatcgtgctgtcacataccaattcagttgttaacccatttatctatgcctatcgcatccgggagttt
cgccaaacttttaggaagataatcaggagtcatgtcctgcggcagcaggaaccatttaaagcggcagggacatccgcacgcgttttg
gcggcgcatggatcagacggggagcaagtatcattgcgcctcaacgggcacccccctggtgtttgggctaatggatcagccccgca
cccagaacggcggcctaatggttacgccttgggcttggtgagcgggggctccgcccaggagtcacaggggaacactggccttcctg
acgtggaactcttgagtcacgagctgaagggcgtttgcccagagcctcccggactcgatgatccactggctcaagatggggctggcg
tgtcatga

**Nanobody Purification from *E. coli.*** BL21(DE3) *E. coli* were transfected via heat shock with pET26b(+) plasmids encoding nanobodies of interest and grown in medium (Terrific Broth) containing ampicillin (100 µg/mL). Transformed bacteria were used generate a starter culture, which was used to inoculate full-size cultures (1-4 L) containing antibiotic. This culture was grown at 37°C with growth monitored through measurement of the optical density at 600 nm ($OD_{600}$). When $OD_{600}$ values between 0.3 and 0.8 were observed, protein expression was induced by addition of isopropyl β-d-1-thiogalactopyranoside (IPTG, 1 mM). The induced culture was then shaken 30°C overnight.

Bacteria were harvested via centrifugation for 30 min at 6,000 RPM (Avanti J Series centrifuge). Cells were resuspended in 30 mL of NTA wash buffer (tris buffered saline + 10 mM imidazole, pH 7.5) containing protease inhibitor (Pierce Protease Inhibitor Tablets, ThermoFisher A32953). Cells were then lysed via sonication and the lysate was centrifuged at 15,000 RPM for 45 min. The supernatant was then passed through a fritted column containing nickel NTA beads (His Pur$^{TM}$ Ni-NTA Resin) equilibrated with Nickel NTA wash buffer. After initial flowthrough, beads were washed 3x with Nickel NTA wash buffer. Subsequently, bound protein of interest was eluted using 10 mL of Nickel NTA elution buffer (TBS + 150 mM imidazole, pH 7.5). Sample was subjected to size exclusion chromatography (Cytiva Akta$^{TM}$ / Pure) using a HiLoad $^{TM}$ 16/600 Superdex 200 pg column with an isocratic gradient of TBS (flow rate 1 mL/min). Fractions of interest were collected and concentrated via centrifugation using spin filtration columns (Amicon Ultra-15, regenerated cellulose, 10,000 nominal molecular weight limit). Protein concentrations were determined absorption at 280 nm.

**Peptide Synthesis, Cleavage, and Purification.** All peptides were synthesized via Fmoc solid phase peptide synthesis on a Gyros PurePep Chorus Automated Peptide Synthesizer. Peptide assembly was performed on Rink Amide resin (0.05 mmol scale) to afford a C-terminal carboxamide. Fmoc-amino acids were dissolved in dimethylformamide (DMF) and added to resin (8 equivalents) with HATU ((1-[Bis(dimethylamino)methylene]-1H-1,2,3-triazolo[4,5-b]pyridinium 3-oxid hexafluorophosphate, 8 equivalents) and N,N-diisopropylethylamine (DIPEA, 16 equivalents). Fmoc groups were deprotected using 20% piperidine in DMF. For peptides bearing a N-terminal fluorescein, the fluorophore was manually coupled after automated linear synthesis using 10 equivalents of 5(6)-Carboxyfluorescein (Acros Organics), HATU (10 eq), and DIPEA (20 eq) and overnight incubation at room temperature.

Cleavage of peptides was performed using a cleavage cocktail comprised of trifluoroacetic acid(TFA)/$H_2O$/triisopropylsilane(TIS) (92.5:5:2.5% by volume). Variants containing cysteine residues were cleaved using the following cleavage cocktail TFA/$H_2O$/TIS/ethanedithiol (EDT)

(90:5:2.5:2.5 by volume) and rocked at room temperature for 3 hours prior to filtration. Product was precipitated using chilled diethyl ether and pelleted by centrifugation (3,000 RPM for 2 minutes). Diethyl ether was decanted, and the pellet was dried under $N_2$ prior to being dissolved in DMSO and purified by HPLC. Peptides were purified via preparative-scale HPLC using a Phenomenex Aeris Peptide XB-C18 Prep column (particle size 5 µM, 100 Å pore size) with a linear gradient of solvent A (0.1% TFA in $H_2O$) and solvent B (0.1% TFA in acetonitrile). Fractions of interest were combined and lyophilized. Lyophilized peptides are then dissolved in DMSO at desired concentrations and frozen.

**Flow cytometry analysis of peptide binding to HEK293 cells.** HEK293 cells stably expressing A2AR($Nb_{6E}$) were cultured as described above. Cells were harvested by trypsinization, which was quenched with the addition DMEM/FBS. Cells were then transferred to a round bottom 96 well plate, pelleted by centrifugation (500 rpm for 3 min.), and resuspended in PBS containing 2% BSA (w/v) (PBS/BSA), and pelleted a second time. The washed cell pellets were then resuspended in PBS/BSA with 10 nM tracer peptide (FAM-6E-C14) mixed with varying concentrations of unlabeled competitor peptides for 30 minutes. Cells were pelleted, washed, then resuspended in PBS/BSA containing Alexafluor647 conjugated anti-fluorescein antibody (1:1000 dilution in PBS/BSA, (Jackson ImmunoResearch, 200-602-037)) and incubated for 30 minutes on ice prior to washing. Washed cells were then resuspended in PBS/BSA for analysis by flow cytometry on a CytoFlex flow cytometer (Beckman Coulter). Live cells were identified based on forward scatter/side scatter profile and staining intensity was monitored in the Alexafluor647/APC channel. A minimum of 2,000 events corresponding to live cells were recorded. Flow cytometry histograms were used to calculate median fluorescence intensity (MFI) values for the APC channel in each sample. MFI values were averaged among replicate samples.

**Fluorescence microscopy analysis of nanobody staining.** A 4-chamber glass slide (Lab-Tek II Chamber SlideTM) was treated with 10% polylysine solution (500 µL/chamber) for 5 minutes, rinsed with molecular-biology grade water and allowed to dry for 1 hour. Trypsinized HEK293 cells were harvested, transferred to a 4-well glass slide and left to incubate in DMEM/FBS at 37°C overnight at which point a confluency of approximately 90% was reached. The medium was aspirated, cells were washed once with PBS/BSA (1 mL) and placed on ice. Cells were stained with 500 µL of indicated solutions for 30 minutes. After incubation, chambers were subsequently washed with PBS/BSA and then fixed with 4% paraformaldehyde solution for 15 minutes. The chambers were then washed with PBS/BSA. Following the wash, the chamber walls were removed and mounting solution containing DAPI was applied to each chamber (ProLong Glass Antifade Mountant with NucBlue). Chambers were enclosed with a coverslip (Gold Seal Cover Glass 24x50 mm No. 1 1/2) and allowed to cure overnight in the dark.

Stained cells were imaged using a Nikon Eclipse 50i microscope coupled to a Cool Snap ES2 CCD camera (Photometrix). Images were acquired at 20x magnification using various filter channels (DAPI, Fluorescein, and Rhodamine). For visualization of each stain, the following exposure times were used: DAPI (5 sec.), Fluorescein (30 sec.), and Rhodamine (30 sec.) Images were processed with ImageJ (FIJI package).

**Liquid chromatography/mass spectrometry (LC/MS) analysis of peptides and proteins.** Mass spectrometry data was acquired on a Waters Xevo qTOF LC/MS or an Agilent Affinity II 6130 quadrupole LC/MS instrument. Samples were resolved by reverse-phase LC (Hamilton PRP-h5 column, 5 µM particle size, 300 Å pore size) and analyzed in positive ion mode. For proteins analyzed by mass spectrometry singly charged ions were not observed, so protein intact mass was calculated from analysis of multiply charged ions using the MaxENT algorithm on MassLynx software.

**Protein labeling via sortagging.** Sortagging reactions were comprised of the following components: protein bearing a sortase recognition motif (LPETGG) followed by a hexa histidine tag at the C-terminus (20-200 µM final concentration), triglycine-probe conjugates (500-1000 µM final concentration), and Sortase 5M (10-20 µM final concentration). Reactions were performed sortase buffer (10 mM $CaCl_2$, 50 mM Tris, 150 mM NaCl, pH 7.5) and shaken overnight at 12°C overnight. After incubation, the reaction was incubated with nickel NTA beads to capture Sortase 5M and unreacted starting protein. Uncaptured material was further purified using disposable desalting columns to remove triglycine conjugates (Cytiva PD-10 Sephadex™ G-25M). Fractions containing product were combined then concentrated by spin filtration (Amicon Ultra 0.5 mL Centrifugal Filters 10,000 NMWL).

**Surface plasmon resonance (SPR) binding experiments.** SPR measurements were performed on a GE Biacore T100 – T200 Sensitivity Enhanced Instrument using a Cytiva Series S Sensor Chip SA (immobilized streptavidin). Ligand ($Nb_{6E}$-biotin) was prepared using standard sortase ligation protocols described above and diluted to a concentration of 80 µg/mL. The SA chip was conditioned using successive treatments with 1) 1M NaCl, 50 mM NaOH 2) 50% isopropanol, 50 mM NaOH, 1M NaCl.

Analyte samples (6E and analogues) were prepared via two-fold serial dilutions in PBST ranging from 1.6 µM to 3.125 nM. Analyte was flowed over the chip at 50 µL/min with a contact time 60 seconds, and a dissociation time of 120 sec. Regeneration step (to dissociate 6E peptides) included of consecutive washes of 10 mM glycine solution (pH 1.5). Sensorgrams were fitted using the Biacore T200 evaluation software to a 2-state model, with local Rmax. Raw sensorgrams and their respective fits were exported as ASCII files and regraphed on GraphPad prism.

A two-state binding model in which each concentration of peptide was fit to a local maximum signal intensity (two-state model, local Rmax) was used for analysis because of poor concordance between experiment and model when using one-state and consistent Rmax models. The cause of poor modeling with one-state models and the need for local Rmax models is unclear, but it may relate to different conformational states or oligomerization states for the immobilized $Nb_{6E}$.

**Synthetic protocols:**

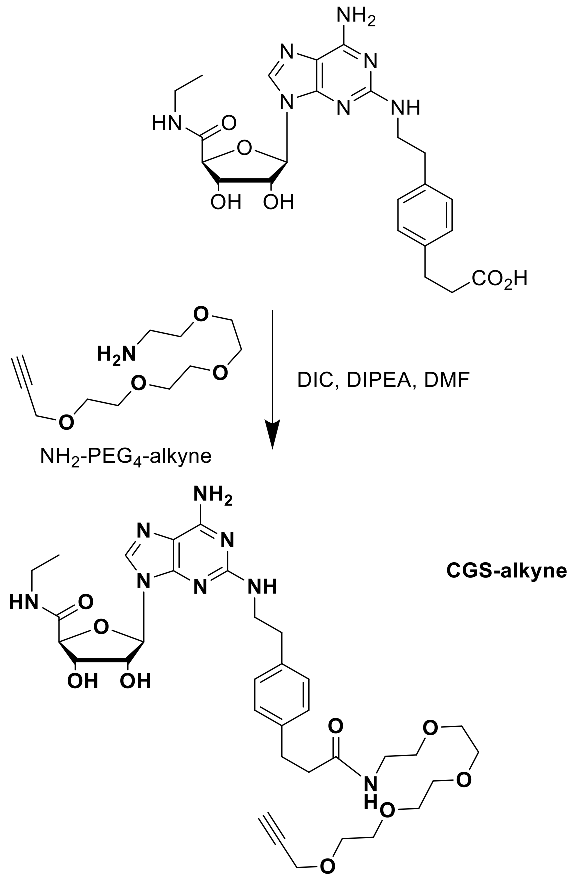


**CGS-alkyne**. CGS21680 (Cayman Chemical, # 17126) was dissolved in dimethylformamide (DMF) and diisopropylcarbodiimide (1 equivalent) was added. To this mixture 2 molar equivalents of $NH_2$-$PEG_4$-alkyne was dissolved in DMF and was added (Click Chemistry Tools, # TA101-100). Next, diisopropylethylamine (5 molar equivalents) was added. The reaction was shaken at 10°C overnight. The reaction was purified by reverse-phase preparatory HPLC (C18 column, gradient 20-90% acetonitrile in water with 0.1% trifluoroacetic acid).

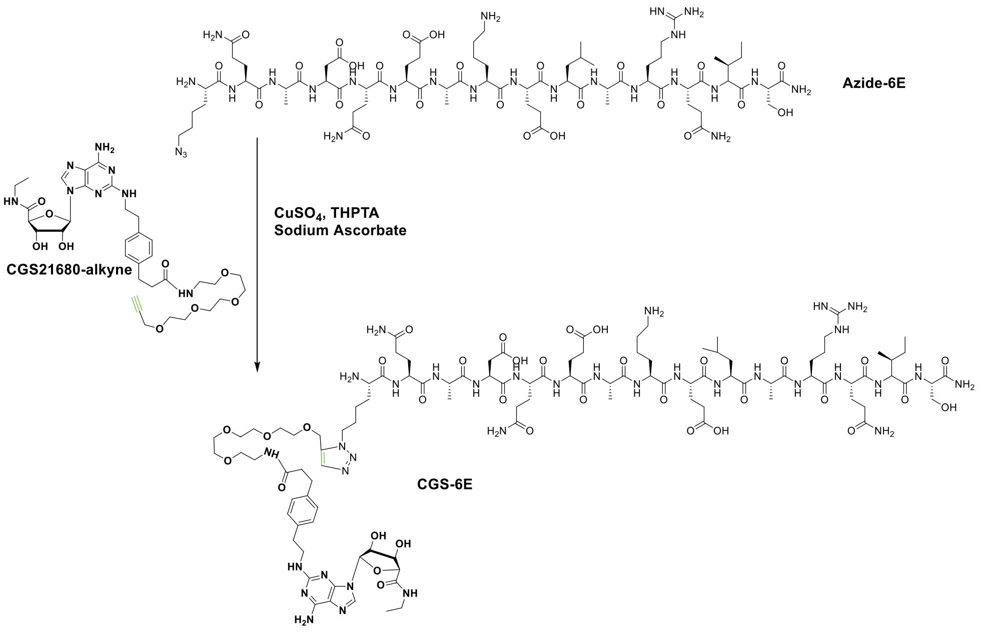


### CGS-6E

### Copper-catalyzed click chemistry (CGS-6E)

Azide-alkyne conjugation was performed as previously described[8] with some modifications. Azide-6E was dissolved in DMF (5 mM) and CGS-alkyne was added at a 3-fold excess (15 mM). $CuSO_4$ heptahydrate was dissolved in water and mixed with THPTA (Click Chemistry Tools, #1010-100) at a 1:5 molar ratio to prepare a 10x stock solutions (1 mM $CuSO_4$, 5 mM THPTA). The pre-mixed copper-THPTA solution was then added to the DMF solution of azide and alkyne. A fresh stock solution of 100 mM sodium ascorbate was prepared in water and diluted 1:20 into the reaction solution to initiate the reaction (final concentration of 5 mM). The reaction was monitored by LC/MS and additional aliquots of premixed $CuSO_4$/THPTA, and sodium ascorbate were added over the course of 12-48 h to promote conversion to product. The reaction was purified by reverse-phase HPLC, lyophilized, and dissolved in DMSO at a concentration of 1 mM for use in biological assays.
